# Astrocytic response to traumatic brain injury to rescue neuronal mitochondrial dysfunction through mitochondrial transfer

**DOI:** 10.64898/2026.01.22.701145

**Authors:** Gopal V. Velmurugan, Hemendra J. Vekaria, Alexander G. Rabchevsky, Kai Saito, Josh M. Morganti, Samir Patel, Brad Hubbard, Patrick G. Sullivan

**Author notes:** Corresponding author: Patrick G. Sullivan, Ph.D., 475 Biomedical & Biological Sciences Research Bldg 741 South Limestone St Lexington, KY, USA 40536-0509. Co-Corresponding author: Velmurugan Gopal Viswanathan, Ph.D. 451 Biomedical & Biological Sciences Research Bldg 741 South Limestone St Lexington, KY, USA 40536-0509.

## Abstract

As highly dynamic organelles, mitochondria play an essential role in neuronal survival and synaptic function. Excitotoxicity is as a critical factor that promotes mitochondrial dysfunction after traumatic brain injury (TBI). Intercellular mitochondrial transfer and exogenous mitochondrial transplantation are emerging concepts to understand mitochondrial trafficking in response to mitochondrial dysfunction; however, robust *in vivo* evidence remains limited on the extent of these processes in the central nervous system (CNS). There is a significant knowledge gap in our understanding of mitochondrial transfer mechanisms under both normal physiological conditions and after experimental TBI. Mouse lines expressing mitochondrial green-fluorescent dendra-2 (mtD2) and GFP (mtGFP) targeted to inner and outer mitochondrial membranes, respectively, were used to study astrocyte-specific (Aldh1l1-CreER; mtD2^f/f^ – AmtD2 and Aldh1l1-CreER; mtGFP^f/f^ – AmtGFP) and neuron-specific (CamK2aCre; mtD2^f/f^ – NmtD2 and CamK2aCre; mtGFP^f/f^ – NmtGFP) mitochondrial dynamics and bioenergetics in acute TBI and excitotoxicity. At 24 hrs following TBI, neurons in the NmtD2 mouse brain exhibited rapid and significant alterations in mitochondrial morphology, including changes in total mitochondrial volume, volume distribution, and sphericity. Synaptic neuronal (SN) mitochondria display robust deficits in mitochondrial bioenergetics and complex protein levels while non-synaptic neuronal (NSN) mitochondria show State III bioenergetics and complex proteins at control levels. These findings are accompanied by a marked increase in astrocyte-derived mitochondria (AmtGFP) transfer to neurons at 24 hrs post-injury, compared to control animals, but no increase in transfer to neuronal synapses. While TBI also altered astrocytic mitochondrial morphology in the cortex, astrocytic mitochondrial bioenergetics remained preserved. Single-cell RNA-seq analysis of astrocytes revealed significant transcriptional reprogramming following TBI, characterized by the upregulation of genes associated with mitochondrial homeostasis and the machinery for organelle trafficking. *In vitro* co-cultures of primary cortical astrocytes and neurons demonstrated that astrocytes can transfer mitochondria to neurons via direct contact and that NMDA-mediated excitotoxicity further enhanced this astrocyte-to-neuron mitochondrial transfer. Furthermore, astrocytic-derived extracellular vesicles containing mitochondria (EV-mito) deliver mitochondria to neurons and EV-mediated mitochondrial transfer significantly ameliorated NMDA-induced mitochondrial dysfunction in primary cortical neurons. Together, these findings show that astrocytes take on a TBI-related phenotype that facilitates dynamic changes in mitochondrial networks and mitochondrial trafficking to neurons. Astrocytic transfer of respiratory-competent mitochondria support is an intrinsic neuroprotective response to injury that supports mitochondrial function in neuronal soma, dendrites, and axons but not at the neuronal synapse. Finally, we show therapeutic potential of exogenous mitochondrial transfer, particularly via EV-mito, for treating neurological disorders associated with excitotoxicity, such as TBI.

## Introduction

Traumatic brain injury (TBI) is a leading cause of disability that affects millions of individuals each year in the United States^1^. People who sustain a TBI often experience a wide range of persistent symptoms, including cognitive impairment, depression, and an elevated risk of developing dementia^2,3^. Currently, there are no FDA-approved therapeutics that directly target the neurobiological consequences of TBI, and a substantial knowledge gap remains in our understanding of both the injury processes and their long-term sequelae^4^.

Mitochondria are central players in neurobiological responses to brain trauma including neuronal resilience. Mitochondrial dysfunction is a well-established hallmark of secondary injury after TBI ^5–9^. One critical function of mitochondria is buffering excess calcium during glutamate-induced excitotoxicity, a process that particularly affects neurons^10^. Following trauma, elevations in cytosolic calcium can overload mitochondria and trigger the mitochondrial permeability transition (mPT)^11^. Once initiated, mPT causes mitochondria to release calcium back into the cytosol, setting off a cascade of permeability transition events that culminate in neuronal cell death^12^. One potential protective strategy is to increase the mitochondrial pool within neurons. A larger mitochondrial network increases the cell’s overall calcium buffering capacity, allowing the mitochondria to absorb and store more calcium before any single mitochondrion reaches the threshold for mPT pore opening. Intercellular horizontal mitochondrial transfer (IMT) is a process through which one cell donates mitochondria to another^13,14^, thereby representing a way to expand the neuronal mitochondrial pool in response to cellular insults such as TBI.

IMT has been documented in several organ systems to eliminate defective mitochondria^15,16^ and act as a signaling mechanism for cellular protection^17,18^. IMT also occurs within the central nervous system (CNS) as a means of transmitophagy^19^ and mitochondrial protection after injury^20^. Astrocytes are critical in the acute phase following TBI^21^. Astrocytes are thought to be the primary donor cell type of mitochondria in the CNS, supplying them to both neurons and endothelial cells^20,22–24^. However, it remains undetermined whether astrocytes donate mitochondria to neurons following TBI and how this exchange influences neuronal and astrocytic functions.

The mechanisms of IMT are still debated, involving both contact-dependent routes, such as tunneling nanotubes, and contact-independent pathways, including extracellular vesicles (EV-mito)^25–27^. Astrocytes have also been shown to release mitochondria packaged within EV (Ast-EV-mito)^22^, raising the possibility that this process contributes to endogenous neuroprotection that could be harnessed therapeutically through exogenous mitochondrial transplantation (EMT). Here, we tested the hypothesis that astrocytes transfer mitochondria to neurons after TBI to restore neuronal bioenergetics and function. Using mitochondrial reporter mouse models (mtD2 and targeted GFP), we examined astrocytic and neuronal mitochondria dynamics *in vivo* and *in vitro* under physiological and injury conditions. Specifically, we examined the acute changes in mitochondria shape, volume and function of both neurons and astrocytes following experimental TBI. While characterizing the acute astrocytic responses to injury, we visualized astrocytic mitochondria within both neurons and isolated synaptosomes. Finally, we investigated the EMT response under conditions of excitotoxicity. Together, these studies define astrocytic mitochondrial transfer as a potential protective mechanism in TBI and provide insight into its therapeutic relevance in other CNS disease states.

## Methods

### Experimental animals

All animal experiments were performed in accordance with the NIH Guide for the Care and Use of Laboratory Animals and were approved by the Institutional Animal Care and Use Committee at the University of Kentucky. All the breeder mice were purchased from Jackson Laboratories, bred and maintained at the Division of Laboratory Animal Resources (DLAR) at the University of Kentucky. Throughout the study, animals were socially housed in individually ventilated cages on a 12 hrs light cycle and had ad libitum access to food and water. *Global mtD2 mice*: Mice (B6;129S-Gt (ROSA)26Sortm1.1 (CAG-COX8A/Dendra2) Dcc/J) expressing a mitochondrial-specific version of Dendra2 green (mtD2) was used to globally express mtD2 (Strain #018397). *Astrocytic mitochondrial reporter mice:* A tamoxifen inducible astrocyte-specific mitochondrial reporter mouse expressing mtD2 (AmtD2) targeted to the inner mitochondrial membrane (IMM) and GFP (AmtGFP) targeted to the outer mitochondrial membrane (OMM) were generated by crossing Mice B6;129S-Gt (ROSA)26Sortm1(CAG-COX8A/Dendra2) Dcc/J (Jax#018385) or B6N.Cg-Gt (ROSA)26Sortm1(CAG-EGFP/Synj2bp) Thm/J (Jax#032675) with B6N.FVB-Tg (Aldh1l1-cre/ERT2)1Khakh/J (Jax #031008), respectively. Cre-positive animals were injected with tamoxifen (TAM) at 75mg/kg body weight around 6-weeks of age for five consecutive days. One month after TAM injection, animals were used for experimental purposes. *Neuronal mitochondrial reporter mice:* Like astrocyte mitochondrial reporter mice, neuron-specific mitochondrial reporter mice were generated by crossing Jax#018385 (mtD2) and Jax#032675 (mtGFP) mice with B6.Cg-Tg(Camk2a-cre)T29-1Stl/J (Jax#005359). All animals were older than 3 months of age because CamK2a expression starts only around 2 months of age.

### Controlled cortical impact

Controlled cortical impact (CCI) was performed as previously described (1.0 mm depth of contusion at 3.5 m/s with a dwell time of 500 ms)^28–30^. In brief, animals were anaesthetized using isoflurane (2-5%), shaved, cleaned, and prepared for the surgical procedure. A Kopf stereotaxic frame was used to generate brain injury under a pneumatic impactor (Precision Science Instruments). A longitudinal skin incision was made down the middle of the head dorsum before craniotomy (4mm) was performed on the left side of the skull between lambda and bregma. Mice were injured by hitting the dura mater of the brain using a 3mm flat-tip impactor. The injury area was cleaned using cotton tips to mitigate any bleeding, the craniotomy was covered with surgiseal, and the wound was closed using staples.

### Immunofluorescence

Brain cryosection (all the mtGFP), paraffin sections (all the mtD2), and cultured primary cells were immunostained as described previously ^30^. 10% Neutral buffered formalin (NBF)-fixed cultured cells and/or brain sections were permeabilized and blocked in phosphate-buffered saline-Tween 20 (PBST) + 0.2% Triton X-100 in + 1% BSA and 10% normal horse serum for one hour at RT. Then, the cells/sections were incubated overnight at 4°C with primary antibody (1:250 dilution; Table 1) in antibody dilution buffer (blocking buffer and PBST at a 1:1 ratio). This was followed by incubation with fluorophore-labelled secondary antibody (1:500 dilution; Table 1) at RT for 1 h. After washing, the cell cultures and/or sections were mounted on glass slides using Vectashield antifade mounting medium with or without DAPI (H-1500 and H-1400; Vector Laboratories, USA). For cells cultured on glass-bottom plates, cells were directly fixed, stained, and imaged without mounting.

### Confocal imaging and image processing

Fluorescence Z-stack images were acquired using confocal microscopes (Nikon A1R and AXR) equipped with 20× air and 100× oil immersion objectives. Image acquisition and processing were performed using NIS-Elements software (version 5.30.05). Z-stack images were denoised and, in some cases, deconvoluted to enhance resolution. As previously reported ^30^, 3D reconstructions were generated from Z-stack data using either NIS-Elements or Imaris software (version 10.2.0), to visualize mitochondrial transfer events between cells. For quantitative analysis, a custom pipeline was created using the General Analysis 3 (GA3) module in NIS-Elements (Figure S3). This pipeline was employed to determine the total surface area of primary neurons and to quantify the number of astrocyte-derived mitochondria transferred into neurons. Neurons were defined as the region of interest (ROI) within 3D reconstructions for analyses.

### Primary neuron and astrocyte-neuron co-culture

Brain cells were isolated via enzymatic digestion as previously described, with slight modifications^22^. Astrocyte-neuron co-cultures were established using a 1:1 ratio of astrocyte and neuronal culture media. Astrocyte medium was obtained from ScienceCell (MSPP-1801) and neuronal medium consisted of Neurobasal-A (Gibco; 10888022) supplemented with B-27 (Gibco; A3582801) and Penicillin-Streptomycin-Glutamine (Fisher Scientific; 10378016). Cells were cultured on poly-D-lysine-coated glass-bottom 24-well plates (Fisher Scientific; A3890401; VWR; 82050898). Briefly, six P0 AmtD2 pups from the same litter were used. After removal of the meninges, cortical tissues were dissected, minced, and digested in 0.05% trypsin for 15–20 minutes at 37 °C with gentle agitation every 5 minutes. Digestion was halted using DMEM supplemented with 10% fetal bovine serum (FBS), followed by gentle trituration (∼15–20 passes) to dissociate the tissue. The suspension was allowed to settle for 1 minute to remove debris, and the supernatant was then passed through a 70 µm cell strainer. Cells were pelleted by centrifugation at 300 × g for 5 minutes at room temperature and resuspended in high-glucose DMEM supplemented with 10% FBS and antibiotics (ThermoFisher; 15-140-122). Cells were plated and maintained in the initial culture medium for 12 hours, after which the medium was replaced with astrocyte-neuron co-culture medium. Media changes were performed every 3 days. At 4 days in vitro (DIV4), cells were treated with 2 μM 4-hydroxytamoxifen (4-HT; Cayman, 14854) for two consecutive days to induce Cre-mediated recombination. At DIV8, genotyping revealed that 3 out of 6 pups were positive for Aldh1l1-CreER, resulting in mtD2 expression in astrocytes. And these Cre-positive cells were either treated with NMDA (100 µM) for 12 hrs or left untreated as controls. Following treatment, cells were fixed for immunostaining and imaging.

### Primary cortical neuron culture

Primary cortical neurons were isolated from wild-type mouse embryos at embryonic days 13–15, following the same dissociation and plating protocol described above, with the exception that only Neurobasal medium supplemented with B-27 and GlutaMAX was used for neuronal culture. Cells were cultured either in 24-well glass-bottom plates for immunostaining or in 96-well Seahorse XF culture plates for mitochondrial stress test (MST) assays at DIV 8-13.

### Primary astrocyte culture and astrocytic EV-mito enrichment

Primary cortical astrocytes cultures were derived from 1- to 3-day-old postnatal global mtD2-expressing pups, following the same digestion protocol as described above, with the exception that only astrocyte culture medium with supplements was used for astrocyte culture. Upon reaching confluency (8-10 days), primary astrocytes were isolated by first removing nonadherent glial cells through orbital shaking at 200 rpm for 6 hours. Then, the astrocytes were dissociated using trypsinization and sub-cultured for further experiments. After four days, primary astrocyte conditioned media was centrifuged at 13,000 g for 30 minutes at 4 ℃ to enrich astrocytic EV-mito.

### Tetramethylrhodamine, ethyl ester (TMRE) staining

Isolated mitochondria or EV-derived mitochondria were incubated with 500 nM TMRE in respiration buffer containing pyruvate, malate, and ADP for 10 minutes at room temperature. Excess dye was removed by washing prior to imaging.

### Astrocytic EV-mito treatment and Mitochondrial stress test (MST)

Primary cortical neurons (25-30,000 cells/well) were treated with or without NMDA (100 µM) for 12 hours. Cells were then washed and subsequently incubated for 24 hrs with astrocyte-derived EV-mito (Ast-EV-mito) isolated from the conditioned media collected from five confluent T-75 flasks of astrocyte cultures before isolation and pelleting by centrifugation, as described previously. The resulting EV-mito pellet was resuspended and used to treat 18 wells of primary neurons seeded in a 96-well Seahorse XF cell culture microplate. Using a Seahorse XFe96 Flux Analyzer (Agilent Technologies, Palo Alto, CA, USA), MST was performed according to the manufacturer’s instructions. Briefly, the cells were incubated in XF assay medium supplemented with substrates (1 mM pyruvate and 2 mM L-glutamine) for 1 hr before the oxygen consumption rate (OCR) measurement. After three measurements of baseline OCR, respiratory chain inhibitors/uncouplers were added sequentially into each well as follows: 1 µM Oligomycin, 4 µM FCCP, and 0.5 µM Rotenone/Antimycin. After each injection, an additional three OCR readings were taken. Different OCR parameters were calculated by the Wave software version (Agilent Technologies). Final OCR measurements were normalized to the number of cells. The Experiment was repeated 2 times with 5-10 technical replicates for each treatment condition.

### Western blot

Western blot analysis was performed for mitochondrial OXPHOS complex proteins. Protein lysate was prepared from isolated mitochondrial fractions using RIPA buffer (150 mM NaCl, 1% Triton X-100, 0.5% sodium deoxycholate, 0.1% SDS, 50 mM Tris, pH 8.0) and centrifuged at 16,100× g for 30 min. The total protein levels were estimated from the supernatant using a BCA kit (23225, Thermofisher). Western blot samples were obtained using XT sample buffer (1610791, Biorad, Hercules, CA, USA) with DTT and boiled at 95 °C for 10 min. The samples (6 µg protein) were resolved in a 4–12% BIS-TRIS gel (3450125, Bio-Rad, Hercules, CA, USA) under reducing conditions and transferred to a nitrocellulose membrane. Probing was performed using a total OXPHOS antibody cocktail (1:1000; ab110413, Abcam). The signals were detected using IRDy 68RD goat anti-mouse (1: 10,000; 926-68070, Li-Cor, Lincoln, NE, USA). The protein levels were quantified with densitometric analysis of the Western blot bands using ImageJ software.

### Cell-specific fractionated mitochondrial magnetic separation

#### Neuronal mitochondrial fractions

Purified neuronal mitochondrial fractions were obtained from NmtGFP mice using magnetic bead-based immunoisolation with modifications to previously described protocols^31^. Briefly, brain tissue (cortex, hippocampus, or pooled regions) was homogenized in mitochondrial isolation buffer (IB) using a Dounce homogenizer (8–10 strokes). Homogenates were centrifuged at 1,300 × g for 3 min at 4 °C, and the supernatant was collected. At this stage, small aliquots were reserved as the total mitochondrial fraction for use as controls. The remaining supernatant was incubated with anti-GFP antibody (25 µL per 20 mg tissue) in a total volume of 10 mL per LS MACS column (Miltenyi). Following 20 min of rotation at 4 °C, the suspension was applied to the column. The flow-through (elute) contained synaptosomes, whereas the retained fraction represented the somatic mitochondrial pool that was recovered by flushing the column with 1.5 mL IB, followed by centrifugation (13,000 × g, 10 min, 4 °C). Pellets were resuspended in IB, protein concentrations quantified, and aliquots prepared for downstream assays. The synaptosome-containing elute was subsequently centrifuged (13,000 × g, 10 min, 4 °C) and pellets were resuspended in 500 µL IB. Synaptosomes were disrupted using a nitrogen bomb (1,200 psi, 10 min) to release synaptic mitochondria. The lysate was then incubated with anti-GFP antibody–conjugated beads (20 min, rotation), passed through a fresh LS column, and separated into two fractions: (i) non-neuronal (NN) mitochondria, recovered from the column flow-through and (ii) synaptic mitochondria, eluted by flushing the column with 1.5 mL IB. Both synaptic and NN fractions were centrifuged at 13,000 × g for 10 min at 4 °C, pellets were resuspended in IB, protein quantified and used for downstream applications.

#### Astrocytic mitochondrial fractions

Astrocyte-specific mitochondria were isolated from AmtGFP mice using a similar column-based approach. Following brain homogenization and MACS column binding, astrocytic mitochondria were recovered by flushing the column, like the soma fraction and used for downstream application after quantification. Synaptosomal fractions were collected in parallel, pelleted by centrifugation, resuspended in IB, and processed for immunostaining on glass slides to confirm mitochondrial transfer from astrocytes to synaptosomes.

### Mitochondrial bioenergetics measurement

Following cell type–specific mitochondrial isolation, bioenergetic function was assessed using the Seahorse XFe96 Analyzer (Agilent Technologies, Santa Clara, CA, USA) as previously described ^28,31,32^. Mitochondrial suspensions were diluted in respiration buffer consisting of 125 mM KCl, 0.1% fatty acid–free BSA, 20 mM HEPES, 2 mM MgCl₂, and 2.5 mM KH₂PO₄, adjusted to pH 7.2 with KOH. Mitochondrial protein was loaded into Seahorse assay plates at 2.5 µg/well for soma fractions and 4 µg/well for total, synaptic, non-neuronal (NN), and astrocytic fractions (3–5 technical replicates per sample). OCR was recorded following sequential injections of substrates, inhibitors, and uncouplers of the mitochondrial electron transport chain (ETC), prepared in respiration buffer lacking BSA. *State III respiration* (ATP-linked respiration): Pyruvate (5 mM) and malate (2.5 mM) were added as Complex I-specific substrates, followed by ADP (4.3 mM) to stimulate ATP synthase. *State IV respiration* (proton leak respiration): Oligomycin (2.5 µM) was added to inhibit ATP synthase. *State V(CI) respiration* (Complex I–driven uncoupled respiration): FCCP (4 µM), a protonophore uncoupler, was added to collapse the proton gradient and maximize respiration. *State V(CII) respiration* (Complex II–driven uncoupled respiration): Rotenone (0.8 µM) was added to inhibit Complex I, followed by succinate (10 mM) as a Complex II-specific substrate.

### Brain tissue harvesting for single-cell transcriptomics

At the prescribed interval, mice (n = 2 per group) were anesthetized with 5.0% isoflurane before exsanguination and transcardial perfusion with ice-cold Dulbecco’s phosphate-buffered saline (DPBS; Gibco #14040133). Following perfusion, brain tissues were removed, and the pericontusional cortex or analogous region in sham mice was rapidly dissected.

#### Single cell sequencing tissue prep

This dissected tissue from each mouse was immediately transferred into a gentleMACS C-tube (Miltenyi #130-093-237) containing Adult Brain Dissociation Kit (ADBK) enzymatic digest reagents (Miltenyi #130 107 677) prepared according to the manufacturer’s protocol. Tissues were dissociated using the “37C_ABDK” protocol on the gentleMACS Octo Dissociator instrument (Miltenyi #130-095-937) with heaters attached. After tissue digestion, cell suspensions were processed for debris removal following the manufacturer’s suggested ABDK protocol. Following completion of this protocol, cell suspensions were pelleted at 300x*g* for 3 min and gently resuspended in 200 μL of DPBS + 0.4% bovine serum albumin (Invitrogen #AM2616). Cells were sequentially filtered two more times using Flowmi cell strainers (70 μm pore size, Bel-Art #H13680-0040). Cell counts and viability were assessed using AO/PI cell viability dyes (Logos Biosystems #F23001) in tandem with the CellDrop automated cell counter (DeNovix). All samples had >90% viable cells and were diluted according to 10x Genomics’ suggested concentrations for capturing approximately 20k cells per library (Next GEM 3’ Gene Expression v3).

#### Processing of single-cell FASTQ files, dimension reduction, and cell clustering

Pre-processing of scRNAseq data was accomplished using Cell Ranger (v8.0.0, 10x Genomics), with Illumina files aligned via STAR (2.5.1b) to the genomic sequence (introns and exons) using the Mouse (GRCm39) 2024-A annotation. Standard pre-processing protocols were followed in Cell Ranger to identify cells above background and the resulting filtered gene matrices were utilized in Seurat (v5) for single-cell analyses. Genes that were detected in 3 or more cells were used for analyses. Single-cell QC was performed to exclude cells with less than 200 genes or those that exceeded 2500 genes, and greater than 25% mitochondrial genes. Following QC, the two sample objects were merged into a single Seurat object. Data were normalized, scaled, and variable features found using *SCTransform* with default parameters for the merged object. Subsequently, PCA and UMAP were used for two-dimensional visualization of the dimensionally reduced dataset (with the top 30 PCs used, based upon the total variance explained by each). *FindMarkers* function was used to identify the top10 biomarkers for each cell cluster. Cell clusters expressing canonical astrocyte-specific genes (i.e. *Aldoc, Slc2a1, Aqp4)* were identified. These cell clusters were subsequently subsetted and stored as a new object, to which the *SCTransform* and dimension reduction, visualization, and cell identification methods described above were re-applied. After subsetting and data normalization, small groups of distinct clusters with differentially expressed genes (DEGs) corresponding to microglia, ependymal cells, and an unclassifiable group were distinguishable from the main astrocyte groups. Following the removal of these unidentified cells, the remaining astrocyte-selected cells were used for the following downstream analyses, described below.

#### Cell type proportion analysis

Cell type proportions were calculated from single-cell metadata by aggregating cell counts by surgical condition (Sham vs CCI) and cluster identity. Pseudocounts of 0.5 were added to avoid zero-inflation issues in compositional data analysis. Bayesian compositional analysis was performed using a Dirichlet regression model implemented in brms v2.22.0 with a logit link function and QR decomposition for numerical stability. MCMC sampling was conducted using 4 chains with 2,000 iterations each. Statistical differences between conditions were assessed using posterior predictions to calculate condition differences (CCI - Sham) for each cell type. Cell types were considered significantly different if the 95% credible interval excluded zero.

#### Mitochondrial pathway enrichment analysis

Single-cell gene set enrichment analysis was performed using escape v2.2.3 to evaluate mitochondrial pathway activity across conditions. Mitochondrial gene sets were obtained from MSigDB collections using msigdbr for Mus musculus. Enrichment analysis was conducted using single-sample gene set enrichment analysis (ssGSEA) with normalize = TRUE and a minimum gene set size of 4 genes. Results were visualized using hierarchical clustering of both rows and columns with scaled enrichment scores.

#### Differential expression analysis

Differential expression analysis was performed between the CCI and Sham conditions using Seurat’s FindMarkers function with the SCT assay and the default Benjamini-Hochberg multiple testing correction. The Wilcoxon rank-sum test was employed with logfc.threshold = 0 and min.pct = 0 to include all genes for comprehensive analysis. Genes were classified as significantly differentially expressed based on adjusted p-value < 0.001 and absolute log2 fold change > 1. MA plots were generated to visualize the relationship between average expression and log2 fold change.

#### Data visualization

Cell type distributions were visualized using stacked bar plots implemented in dittoSeq v1.18.0, displaying proportional representation across experimental conditions and individual samples. Gene expression patterns across clusters were visualized using stacked violin plots implemented in scCustomize v3.1.3.

##### Statistics

Statistical analysis was performed using Graph Pad Prism (GraphPad Software, CA, USA). A significant difference among groups was defined as p < 0.05 for all analyses. The Shapiro-Wilk test was completed to ensure normality. As these criteria were met for all experimental data, parametric statistics were employed for all analyses. A two-way ANOVA with Sidak post-hoc multiple comparisons test or one-way ANOVA with Tukey post-hoc multiple comparisons or unpaired t-test used depending upon the groups.

## Results

### Changes in neuronal mitochondrial morphological dynamics impair synaptic mitochondrial function preferentially over soma after TBI

Changes in mitochondrial dynamics and altered expression of fusion and fission proteins are observed in several neurodegenerative diseases^32,33^. Depending on the context and disease condition, it could be compensatory or detrimental^34–39^. We reported that TBI drastically alters mitochondrial dynamics (size, shape, and count) using mice expressing mtD2 globally^29^. Decades of work from our lab and others have demonstrated that glutamate-mediated excitotoxicity after TBI causes mitochondrial dysfunction through excessive Ca^2+^ influx into neurons^28,31,40–43^. However, the specific effects of TBI on neuronal mitochondrial dynamics and bioenergetics remain undetermined. To address this, we generated two neuron-specific mitochondrial reporter mouse models: 1. NmtD2, which expresses mtD2 targeted to the inner mitochondrial membrane (Fig. S1A), has better fluorescence, suitable for most of the mitochondrial morphological dynamics studies. 2. NmtGFP, which expresses GFP targeted to the outer mitochondrial membrane (Fig. S2A), is suitable for mitochondrial isolation. We validated neuron-specific expression of both reporters by immunostaining brain sections with neuronal markers (NeuN, MAP2 or β3-tubulin) and the astrocytic marker GFAP (Fig. S1B, S2). Change in mitochondrial dynamics was determined in the penumbral region of injured and sham brain sections, both in cortex and hippocampus (Fig. S1C), 24 hrs post-TBI in NmtD2 mice. The percent mitochondrial volume distribution (Fig S1D, G) in both the cortex and hippocampus demonstrated a left shift compared to the sham. Consistently, mitochondrial volume (Fig. S1E, H) significantly decreased while mitochondrial sphericity (Fig. S1F, I) increased dramatically after CCI compared to sham. Since we saw a drastic change in mitochondrial morphological dynamics, we were curious whether they were compensatory or detrimental. To determine this, we isolated neuronal soma, synaptic (Syn) and non-neuronal (NN) fractions of mitochondria using magnetic anti-GFP antibody beads along with total mitochondria from NmtGFP mice (Fig. 1A, B, C). The amount of anti-GFP antibody required per gram of tissue, the concentration of necessary mitochondria per well in the Seahorse XF96 plate, the concentration of complex inhibitor (oligomycin) and uncoupler (FCCP) were optimized, and intact mitochondrial membrane potential was tested before running mitochondrial bioenergetics (Fig. S3A). Results demonstrated that mitochondria from soma (2.5 µg/well) respired at a higher rate compared to mitochondria from the synaptic (4 µg/well) fraction (Fig. 1D, F). Overall, injury decreased OCR at different states of mitochondrial bioenergetics from all four fractions (soma, Syn, NN, and total); (Fig. 1E, G, and Fig. S4A, B). There was a significant OCR decrease only at state V(C-I) of the soma fraction (Fig. 1E) compared to all states in the synaptic fraction (Fig. 1G). When comparing only the injured animals with all four different fractions, the synaptic fraction was most affected with the lowest OCR whereas the soma fraction was least affected with the highest OCR (Fig. 1H). Such functional and dynamic changes in the mitochondria were paralleled by changes in mitochondrial complex protein expression in Western blots (Fig. 1I and K). Western blot quantification from soma (Fig. 1J) and synaptic (Fig. 1L) fractions shows a significant decrease in complex-I, II and V in the synaptic fraction but not in the soma fraction.

**Figure 1.**
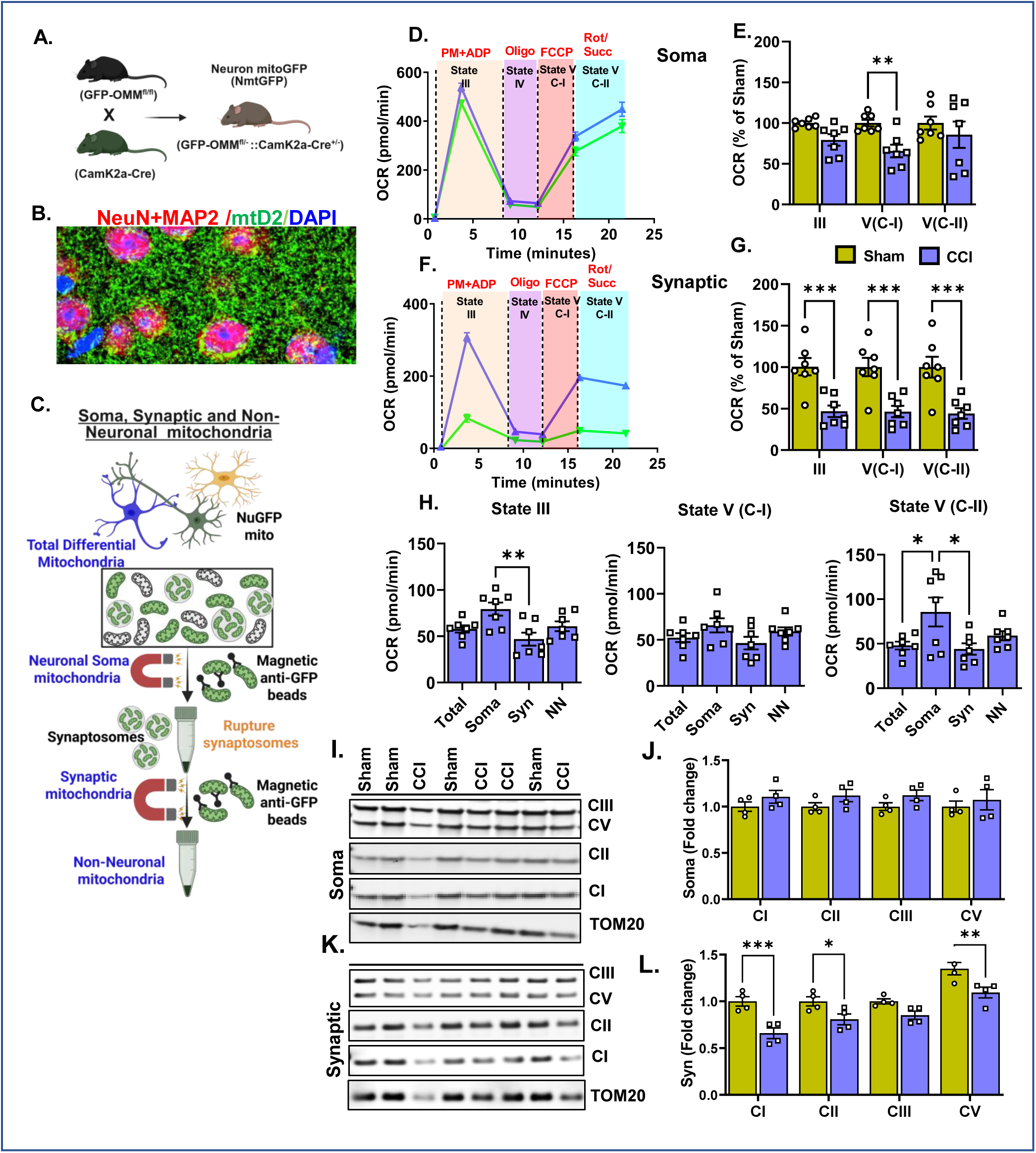
TBI preferentially impairs synaptic mitochondrial function over soma and non-neuronal mitochondrial fraction. **(A)** Schematic illustrating the generation of neuron-specific mitochondrial reporter mice (mtGFP^f/f^; CamK2a-Cre^+/−^) expressing the green fluorescent protein GFP (NmtGFP) targeted to the outer mitochondrial membrane. (B) Representative confocal micrograph of a brain section from NmtGFP mice cortex, stained for MAP2+NeuN (red) and DAPI (blue). (C) Flow diagram outlining the isolation of neuronal soma, synaptic, and non-neuronal (NN) mitochondrial fractions from the brain using anti-GFP magnetic beads in NmtGFP mice, 24 hrs post-CCI. (D, E) Representative traces and quantification of OCR from the neuronal soma fraction of mitochondria isolated from sham and CCI (ipsilateral punch) from two cohorts. Each circle/square represents one animal (n = 7 mice/group). (F, G) Representative traces and quantification of OCR from the neuronal synaptic fraction of mitochondria isolated from sham and CCI. Each circle/square represents one animal (n = 7 mice/group). (H) Bar graphs comparing mitochondrial dysfunction represented as change in OCR (normalized to sham) across four distinct mitochondrial fractions (total, soma, synaptic, and non-neuronal), 24 hrs post-CCI; (n=7 mice/group). (I, J) Representative Western blot micrograph of soma fraction of mitochondria for different mitochondrial respiratory chain complex proteins and quantification of complex I (NDUFSB8), complex II (SDHB), complex III (UQCRC2), and complex V (ATP5A) from one cohort normalized to TOM20I from the same blot. Each circle/square represents one animal (n = 4 mice/group). (K, L) Representative Western blot micrograph of the synaptic fraction of mitochondria for different mitochondrial respiratory chain complex proteins and quantification of complex I (NDUFSB8), complex II (SDHB), complex III (UQCRC2), and complex V (ATP5A), normalized to complex III from the same blot. Each circle/square represents one animal (n = 4 mice/group). Data represent mean ± SEM. P ≤ 0.05 *; P ≤ 0.01 **; P ≤ 0.001 ***; P ≤ 0.001 ***; P ≤ 0.0001 **** by two-way ANOVA with Fisher’s LSD comparison test (E, G, J, L); by one-way ANOVA with Tukey post-hoc test (H).

Together, these results demonstrate that changes in neuronal mitochondrial dynamics following TBI are detrimental, with decreased mitochondrial bioenergetics observed primarily in the synaptic fraction compared to the soma fractions. This observation is supported by a change in mitochondrial complex protein expression. Although synaptic mitochondria participate in the synaptic transmission of action potentials, soma mitochondria appear to respire at a higher rate. This likely maintains neuronal homeostasis and makes them less vulnerable to injury compared to synaptic mitochondria.

### TBI increases mitochondrial transfer from astrocytes to neurons

Astrocytes play a vital role in cerebral metabolism, neurodevelopment, neurotransmission, maintenance of the blood-brain barrier and blood flow^45^. Under physiological and pathological conditions, excess glutamate released from the presynaptic terminal is cleared by astrocytes, which prevents glutamate excitotoxity and neuronal death^46,47^. Also, astrocytes support neuronal viability by transferring functional extracellular mitochondria^20,23,48,49^. However, it is not known whether astrocytes transfer mitochondria to rescue or preserve neuronal function after overt pathology, such as TBI. To address this, we used mitochondrial reporter mice generated in our laboratory (AmtD2 and AmtGFP), and we confirmed cell-specific mitochondrial reporter gene expression in brain sections (Fig. 2A, C and Fig. S5). Naïve mouse brain sections from AmtD2, stained with neuronal marker (NeuN+MAP2) and astrocyte marker (GFAP), were used to determine mitochondrial transfer from astrocytes to neurons under physiological conditions. 3D reconstruction from confocal z-stack images enabled the visualization of astrocytic mitochondria in neurons (Fig. 2B). Importantly, we detected astrocytic mitochondria in proximity to neuronal nuclei using 3D Imaris. This observation eliminates ambiguity of whether they are just juxtapositioned on top of the neurons, not getting inside. To our knowledge, this is the very first time *in vivo* intercellular mitochondrial transfer has been shown at this high resolution in the brain. Next, we examined whether TBI affects astrocyte-to-neuron mitochondrial transfer using AmtGFP mice. We selected this reporter line because it provides superior separation of astrocytic and neuronal mtGFP signals compared to AmtD2 mice, enabling unambiguous and unbiased quantification of transferred mitochondria. Brain sections from the penumbral cortex of injured and naïve mice were stained with neuronal markers (NeuN and MAP2) (Fig. 2D). Using 3D-rendered Imaris reconstructions, we visually detected astrocyte-derived mitochondria within neuronal somata after TBI (Fig. 2G). Defining neurons as the region of interest (ROI), we quantified astrocytic mitochondrial transfer using the unbiased pipeline described in Fig. S6A,B. TBI significantly increased astrocyte-derived mitochondrial transfer to neurons compared to naïve controls (Fig. 2H). To further characterize astrocyte-derived mitochondria at synaptic compartments, we isolated synaptosomes from the cortex of AmtGFP mice (Fig. 2E) and stained them with the synaptic marker synapsin-1 (Fig. 2F). Overall synaptosome abundance was reduced after TBI relative to naïve animals. However, when normalized to total synaptosome number using NIS Elements, the proportion of synaptosomes containing astrocyte-derived mitochondria did not differ significantly between TBI and naïve groups (Fig. 2I).

**Figure 2.**
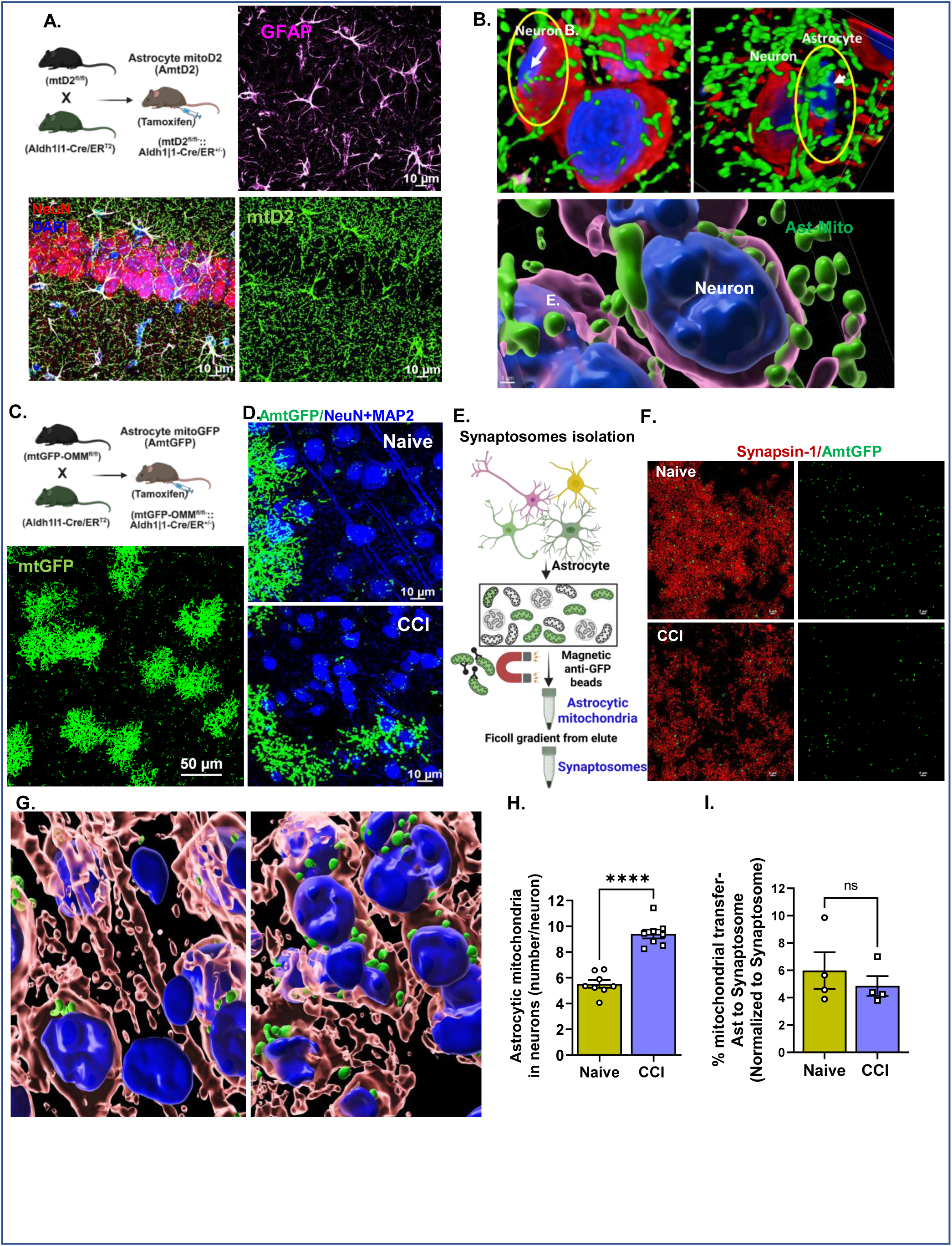
TBI increases mitochondrial transfer from astrocytes to neurons. **(A)** Schematic illustrating the generation of astrocyte-specific mitochondrial reporter mice (AmtD2) and representative confocal micrograph of a hippocampal section from AmtD2 mice. Sections were stained with β3-tubulin + NeuN (neurons-red), GFAP (astrocytes-pink), and DAPI (nuclei-blue), along with astrocyte-expressed mitochondrial reporter signal (mitoD2-green). (B) Representative 3D Imaris reconstruction of a cortical Z-stack image demonstrating astrocyte-derived mitochondria (green) localized within neuronal soma. (C) Schematic showing the generation of the astrocyte-specific mitoGFP reporter line (AmtGFP) and representative confocal micrograph of a cortical section demonstrating selective expression of mitoGFP in astrocytes (green). (D) Immunostained confocal brain sections from naïve and injured AmtGFP mice showing astrocyte-derived mitochondria (green) within neurons (MAP2⁺NeuN-blue). (E) Flow diagram outlining the isolation of synaptosomes from AmtGFP mice using Ficoll gradient centrifugation following depletion of astrocyte-specific mitochondria with anti-GFP magnetic beads. (F) Representative confocal micrographs of isolated synaptosomes from naïve and 24 hrs post-CCI AmtGFP mice. Astrocyte-derived mitochondria (green) within synaptosomes (synapsin-1-red) were quantified using confocal Z-stack imaging and NIS-Elements General Analysis (Nikon). (G) Representative cortical confocal images from naïve and 24 hrs post-CCI AmtGFP mice, with 3D Imaris reconstructions illustrating astrocyte-derived mitochondria (green puncta) within neurons (MAP2+NeuN-blue). Quantification was performed using Z-stack images analyzed in NIS-Elements (Fig. S6 for details). (H) Quantification of astrocyte-derived mitochondria per neuron in naïve and 24 hrs post-CCI mice. Data represent eight random unbiased microscopic fields from 3 naïve mice (139 neurons total) and 4 CCI mice (95 neurons total). (I) Quantification of normalized percent mitochondrial transfer from astrocytes to synaptosomes, calculated by normalizing astrocyte-derived mitochondria to the total synaptosome number using NIS-Elements analysis; n = 4 mice per group. Data represent mean ± SEM. P ≤ 0.0001 **** by unpaired t-test (H, I).

Interestingly, this pattern suggests a correlation between astrocytic mitochondrial transfer and neuronal mitochondrial function. When astrocyte-derived mitochondria did not increase at synapses after TBI, synaptic mitochondrial dysfunction was pronounced. Conversely, in neuronal soma, where mitochondrial transfer was elevated, mitochondrial dysfunction appeared less severe. Together, these findings support the possibility that astrocytic mitochondrial donation acts as a compensatory mechanism to preserve neuronal function following TBI.

### TBI does not affect astrocytic mitochondrial bioenergetics but alters mitochondrial dynamics

Astrocytes play a vital role in cerebral metabolism, neurodevelopment, neurotransmission, maintenance of the blood-brain barrier and blood flow^44^. Under physiological and pathological conditions, excess glutamate released from the presynaptic terminal is cleared by astrocytes, which prevents glutamate excitotoxity and neuronal death^45,46^. Also, astrocytes support neuronal viability by transferring functional mitochondria^20,22,47,48^. However, whether astrocytes engage in mitochondrial transfer to rescue or preserve neuronal function following traumatic brain injury (TBI) remains unknown. To address this gap, we first assessed astrocytic mitochondrial dynamics in the penumbral region of the cortex and hippocampus of AmtD2 mice 24 hrs after injury (Fig. 3A, E). In the cortex of TBI mice, the distribution of mitochondrial volume shifted leftward (Fig. 3B), indicating a fission-like phenotype. Consistent with this, mitochondrial volume decreased significantly (Fig. 3C) while mitochondrial sphericity increased (Fig. 3D), reflecting fragmentation. In contrast, mitochondrial morphology in the hippocampus did not show similar alterations after injury (Fig. 3F–H). Given that changes in mitochondrial dynamics after TBI were associated with reduced neuronal mitochondrial function and that we concurrently observed increased astrocyte-derived mitochondria within neurons, we next asked whether astrocytes transfer respiratory competent mitochondria following injury. To test this, we isolated astrocyte-specific mitochondrial fractions from the cortex and hippocampus of AmtGFP mice (Fig. 3I) and assessed their bioenergetics. Surprisingly, astrocytic mitochondria from both regions maintained normal mitochondrial function 24 hrs after TBI, with no reductions in respiratory capacity compared to naïve controls (Fig. 3J–M). These results suggest that although both neurons and astrocytes respond to TBI by altering mitochondrial dynamics, only neuronal mitochondria become dysfunctional at this early time point, whereas astrocytic mitochondria remain intact. This distinction supports a model in which excitotoxic activation of NMDA and glutamatergic signaling leads to Ca²⁺ overloading and mitochondrial injury in neurons, while simultaneously activating astrocytes. In turn, astrocytes may transfer healthy, respiratory-competent mitochondria to neurons as a compensatory rescue mechanism^50,51^.

**Figure 3.**
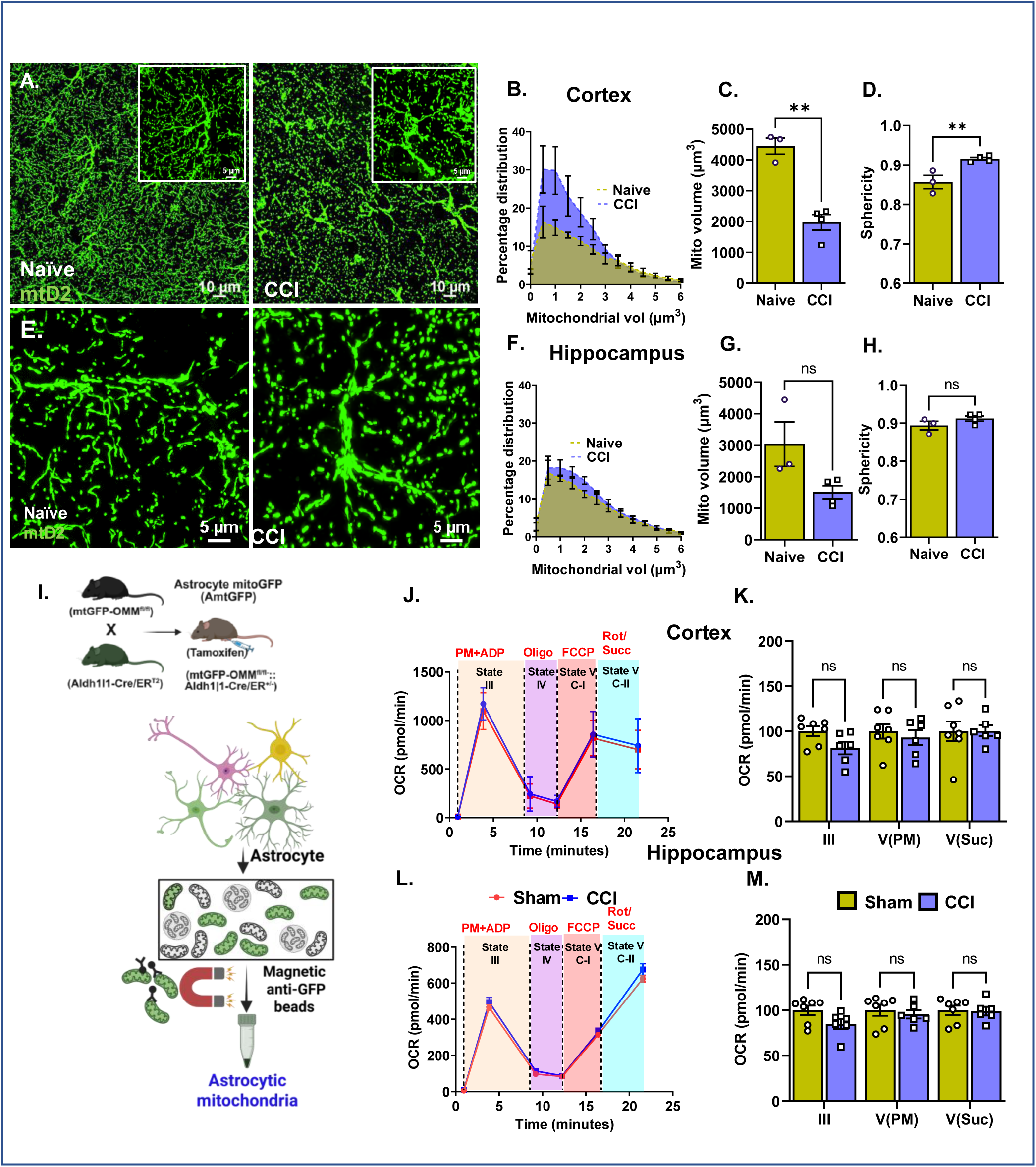
TBI does not affect astrocytic mitochondrial bioenergetics but alters dynamics. (A, E) Representative confocal micrographs of cortical (A) and hippocampal (E) sections from sham and 24 hrs post-CCI AmtD2 mice. (B–D) Quantification of cortical astrocytic mitochondrial percentage distribution (B), mitochondrial volume (C), and mitochondrial sphericity (D), measured from confocal Z-stack images using Imaris. Each circle or square represents one animal (n = 3–4 mice/group). (F–H) Quantification of hippocampal astrocytic mitochondrial percentage distribution (F), volume (G), and sphericity (H), measured from confocal Z-stack images using Imaris. Each circle or square represents one animal (n = 3–4 mice/group). (I) Schematic the workflow of isolating astrocyte-specific mitochondria using anti-GFP magnetic beads from AmtGFP mice. (J, L) Representative OCR traces of isolated astrocytic mitochondrial fractions from cortex (J) and hippocampus (L) from sham and CCI mice (ipsilateral punches). (K, M) Quantification of OCR from astrocytic mitochondrial fractions of cortex (K) and hippocampus (M). Each circle or square represents one animal (n = 6–7 mice/group). Data represent mean ± SME. P > 0.05, non-significant (ns); P ≤ 0.01 ** by unpaired t-test (C, D, G, H); by two-way ANOVA with Fisher’s LSD post hoc comparison (K, M).

### TBI alters gene expression associated with mitochondrial homeostasis and organelle trafficking machineries in reactive astrocytes

Astrocytes became activated within 24 hrs following CCI model of TBI, as demonstrated by immunofluorescence analysis of glial fibrillar acidic protein (GFAP) expression. GFAP staining revealed a significant increase in both expression levels and coverage area in the injured brain compared to sham controls (Fig. 4A, B), corresponding to an increased number of proliferating astrocytes^52^. Reactive astrocytes are known to play a dual role in the pathophysiology of TBI. On one hand, depletion of reactive astrocytes can exacerbate inflammation and promote neuronal death^53^. On the other hand, in transgenic models lacking reactive astrocyte markers such as vimentin and GFAP, neuronal repair mechanisms are enhanced^54,55^. In our experimental conditions, TBI increased astrocyte-to-neuron mitochondrial transfer without altering the functional integrity of the transferred mitochondria. To investigate the underlying mechanisms, we performed single-cell RNA sequencing (scRNA-seq) from the astrocyte population. Consistent with immunohistochemical data, astrocyte-specific transcriptomic analysis revealed eight distinct astrocytic populations, characterized by increased expression of genes associated with astrocyte activation (Fig. 4C, D, E). Pathway enrichment analysis of astrocyte mitochondria revealed that TBI upregulates genes related to mitochondrial biogenesis, oxidative phosphorylation, PGC-1α signaling, mitophagy, extracellular vesicle (EV) biogenesis, and mitochondrial dynamics (fusion and fission) (Fig. 4F). An MA comparing plot of CCI vs Sham showed significant differential expression of key genes involved in EV biogenesis and cargo loading (Sdc4, Cd9, Cd63), EV formation and secretion (Sdcbp), TNT formation and cytoskeletal modelling (Tuba1a, Flna, Fscn1, Actr3), mitochondrial ribosomal (Mrps6, Mrpl33) and mitophagy (Phb2) that were all significantly upregulated in astrocytes 24hrs post-injury (Fig. 5 H). These findings indicate that reactive astrocytes undergo transcriptional reprogramming following TBI to enhance mitochondrial turnover, intercellular communication, and organelle trafficking, thereby equipping them to support neuronal recovery during the acute phase of injury.

**Figure 4:**
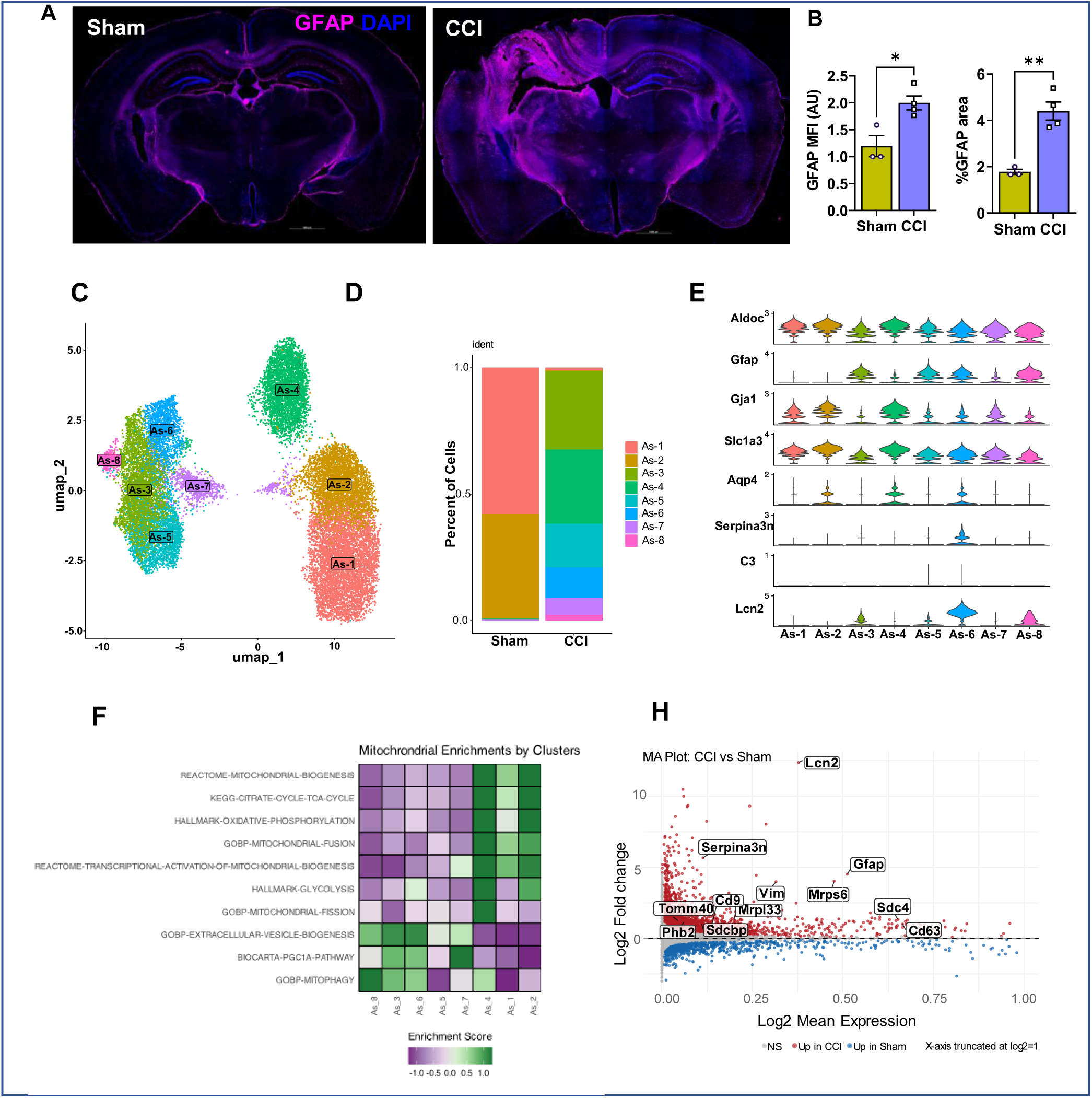
TBI alters gene expression associated with extracellular vesicle biogenesis, trafficking, secretion, and mitochondrial translation in reactive astrocytic population. **(A)** Representative micrographs of whole-brain sections from sham and CCI mice stained with GFAP (astrocytes-pink) and DAPI (nuclei-blue). (B) Quantification of astrocytic mean fluorescence intensity (MFI) and percentage GFAP+ area from brain sections using NIS-Elements General Analysis. Each circle or square represents one animal (n = 3–4 mice/group). (C) UMAP visualization of single-cell transcriptomic profiles from sham and CCI mice (n = 2/group), identifying eight major cell clusters based on canonical marker expression. (D) Proportion of each identified cell cluster represented as a percentage of total cell count in stacked bar plots. (E) Expression patterns of genes associated with reactive astrocytes shown as stacked violin plots across the identified clusters. (F) Astrocyte mitochondrial pathway enrichment analysis displayed as a hierarchically clustered heatmap with scaled enrichment scores. (G) MA plot showing significantly differentially expressed genes, illustrating the relationship between mean gene expression and log2 fold change. Key genes associated with reactive astrocytes, EV biogenesis and cargo loading, tunneling nanotube (TNT) formation and cytoskeletal remodeling, mitochondrial ribosomal proteins, and mitophagy are highlighted.

**Figure 5.**
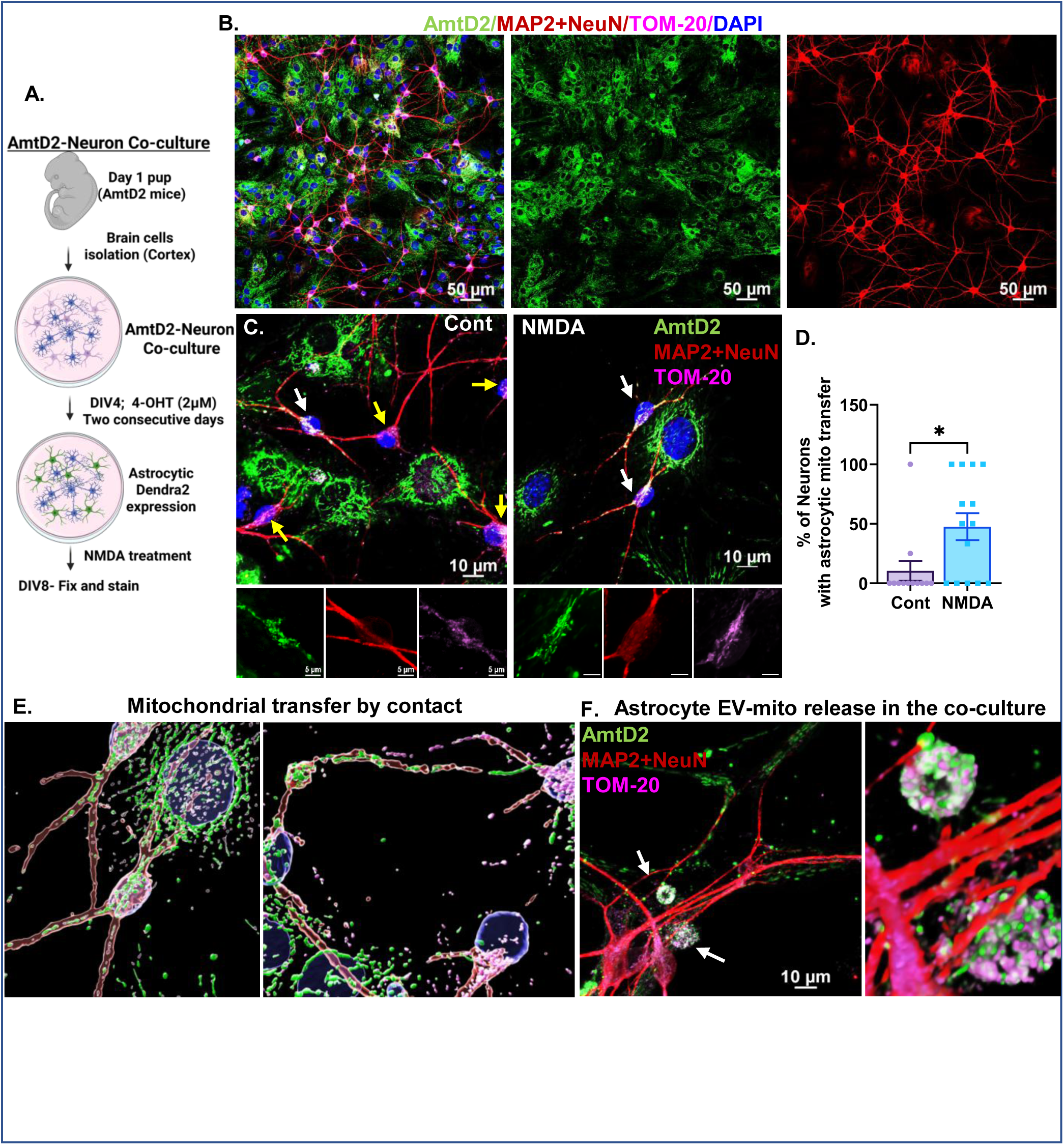
NMDA treatment increases mitochondrial transfer from astrocytes to neurons *in vitro*. **(A)** Flow diagram outlining the brain cell co-cultures from 1-day-old AmtD2 pups and induction using 4-hydroxy tamoxifen (4-OHT) to express specifically mtD2 (green) only in astrocytes to study mitochondrial transfer in neurons. (B) Representative confocal micrograph of brain cell co-culture stained with MAP2+NeuN (neuron-red), TOM20 (mitochondria-green), and DAPI (nucleus-blue), along with 4-OHT-induced mtD2 (astrocyte-green). (C) Representative confocal micrograph of control and NMDA (100µM) treated AmtD2 brain co-cultured cells (DIV-8) stained with MAP2+NeuN (neuron-red), TOM20 (mitochondria-pink), and DAPI (nucleus-blue). Neurons with (yellow arrow) and without (white arrow) astrocytic mitochondria transfer were quantified (D) Bar graph showing the percentage of astrocyte mitochondrial transfer to neurons in the AmtD2 brain co-culture model; n=12-15 fields from 3-Aldhl1l positive pups from the same litter. A total of 17-29 neurons per group was used for the quantification. (E) Representative confocal micrograph of AmtD2 brain co-culture Z-stack 3D images reconstructed using Imaris software showing astrocytic mitochondria (green) transferred to neurons by direct contact from nearby astrocytes. (F) Representative confocal micrograph of AmtD2 brain co-culture showing astrocytic EV-mito (green) labelled with white arrow near the neuronal soma (MAP2+NeuN-red) stained with TOM20 (mitochondria). Data represented as mean ± SME. P ≤ 0.05 * by unpaired t-test (D).

### Excitotoxicity signals increased mitochondrial transfer from astrocytes to neurons in vitro

After TBI, excessive glutamate release leads to excitotoxicity in neurons ^43,56^. At the same time, neuronal glutamate binds to the NMDA receptor, increasing intracellular Ca^2+^ signaling, and activates astrocytes to produce extracellular mitochondria through a CD38- and cyclic ADP-ribose (cADPR)-dependent mechanism^20,57^. Consistent with this framework, our in vivo studies revealed increased astrocyte-derived mitochondrial transfer to neurons after TBI in our transgenic mouse model. To mechanistically test whether NMDA receptor activation enhances mitochondrial transfer, we established an in vitro primary co-culture system using astrocytes and neurons isolated from day 1 AmtD2 pups. Cells were treated with 4-hydroxytamoxifen (4-OHT) at 4 days in vitro (DIV) for two consecutive days, followed by NMDA exposure (Fig. 5A). Within 2 days of 4-OHT treatment, astrocytes robustly expressed mtD2 (Fig. 5B). To distinguish astrocyte-derived mitochondria (green) from endogenous neuronal mitochondria, cultures were stained with the mitochondrial marker TOM20 (pink). At higher magnification, neurons situated in close proximity to AmtD2 astrocytes clearly contained astrocyte-derived green mitochondria (Fig. 5C, white arrows), whereas other neurons did not (Fig. 5C, yellow arrows). Quantification of neurons containing astrocytic mitochondria demonstrated that NMDA treatment significantly increased mitochondrial transfer compared to untreated controls (Fig. 5D). 3D reconstructed images confirmed the presence of astrocyte-derived mitochondria (green) within neuronal soma alongside endogenous TOM20-positive mitochondria (pink) (Fig. 5E). These observations are consistent with previous studies showing mitochondrial transfer via tunneling nanotubes (TNT)^58^. In our earlier work, we demonstrated that astrocytes can also transfer mitochondria through extracellular vesicles containing mitochondria (EV-mito) released from astrocytes to brain capillaries^22^. Similarly, in the current co-culture experiments, we identified astrocyte-derived EV-mito (Ast-EV-mito) adjacent to neurons; these vesicles were positive for mtD2 and co-localized with TOM20 (Fig. 6F).

**Figure 6.**
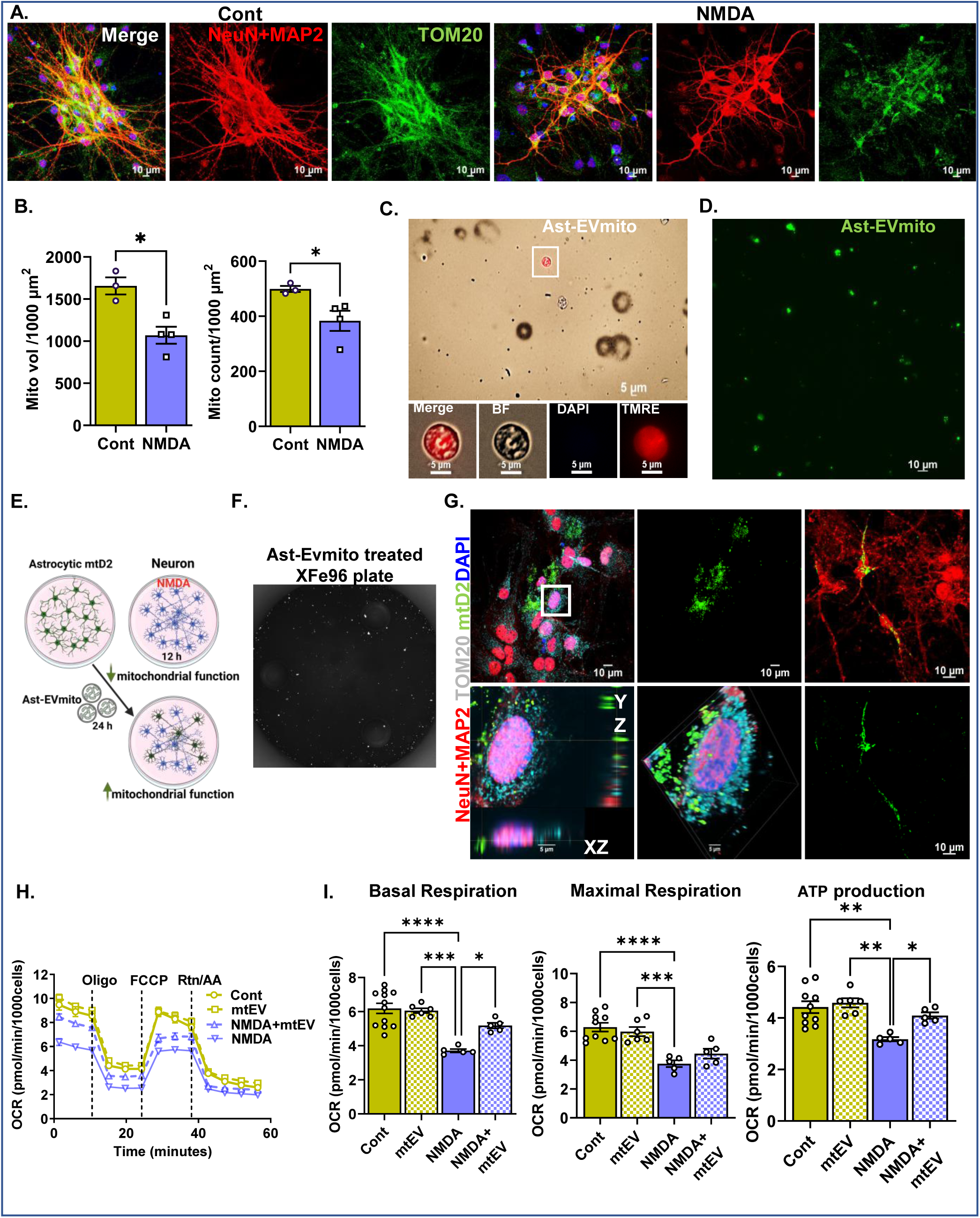
Astrocytic EV-mito improved NMDA mediated excitotoxicity in primary neurons. **(A)** Representative confocal micrographs of control and NMDA-treated (100 µM; 24 hrs) wild-type primary neurons (DIV8) stained with MAP2+NeuN (neurons-red), TOM20 (mitochondria-green), and DAPI (nuclei-blue). Neuronal mitochondrial morphology (volume and count) was quantified using Imaris and normalized to total neuronal surface area using NIS-Elements General Analysis (Nikon). (B) Quantification of mitochondrial volume and mitochondrial count per 1000µm^2^ neuronal surface area in control and NMDA-treated neurons. (C, D) Representative micrographs of isolated astrocyte-derived EV-mito from wild-type (C) and AmtD2 (D) primary astrocyte conditioned media. WT EV-mito were stained with DAPI and TMRE to verify mitochondrial content and confirm the absence of cellular debris. (E, F) Schematic outlining the experimental paradigm for NMDA treatment followed by astrocytic EV-mito administration (E), and the Seahorse XFe96 assay plate well 24 hrs after EV-mito treatment (F). (G) Representative confocal micrograph of primary neurons (DIV8) 24 hrs after treatment with astrocytic EV-mito. Astrocyte-derived mtD2 (green) is visible within neurons (MAP2+NeuN, red), along with endogenous TOM20+ mitochondria (pink), with DAPI-stained nuclei (blue). (H) Representative oxygen consumption rate (OCR) traces from Seahorse mitochondrial stress tests (MST) in neurons treated with Control, EV-mito, NMDA+EV-mito, or NMDA alone. Sequential injections of oligomycin, FCCP, and rotenone/antimycin A are indicated. (I) Quantification of basal respiration, maximal respiration, and ATP production derived from Seahorse XFe96 analysis. Data represent n = 5–12 technical replicates per group, repeated in two independent experiments with similar results. Data represent mean ± SEM. P ≤ 0.05 *; P ≤ 0.01**; P ≤ 0.001***; P ≤ 0.0001**** by unpaired t-test (B); by one-way ANOVA with Tukey’s multiple comparisons test (I).

Taken together, our data suggest that astrocytes can transfer mitochondria to neurons through direct contact-dependent mechanisms as well as through Ast-EV-mito release. Furthermore, NMDA-mediated astrocyte activation significantly enhances the extent of astrocytic mitochondrial transfer to neurons.

### Exogenous Ast-EV-mito treatment improved excitotoxicity-induced mitochondrial dysfunction in primary neurons

Currently, mitochondrial transfer between cells is considered a protective or rescue mechanism ^22,27,58–64^. This phenomenon is being explored as a treatment option through exogenous mitochondrial transplantation for several diseases, notably neurodegenerative diseases^25^. However, several open questions remain in the field of mitochondrial transplantation. What is the fate of transplanted mitochondria? Are they getting integrated along with the host mitochondria?

Do they maintain or improve the mitochondrial function of the host cell? Previous studies, including our own, have demonstrated the feasibility of exogenous mitochondrial transplantation^20,22,65^. However, there is no direct evidence to suggest that excitotoxicity-mediated mitochondrial dysfunction can be rescued by mitochondrial transplantation. In our experiments, primary cortical neurons were treated with NMDA and mitochondrial dynamics (volume and count) were quantified from z-stack confocal images (Fig. 6A). NMDA treatment decreased both mitochondrial volume and number in primary neurons (Fig. 6B). To establish a therapeutic effect, we isolated EV-mito from either WT or AmtD2 primary astrocyte-conditioned media and tested for the presence of mtD2 fluorescence and membrane potential (Fig. 6C, D). NMDA pretreated primary cortical neurons then transplanted with Ast-EV-mito rescued neuronal function (Fig. 6E, F). We were able to visualize transplanted astrocyte-derived green mitochondria juxtaposed to endogenous mitochondria (pink) stained with TOM20 (Fig. 6G). The rescue effect of NMDA treatment on neuronal dysfunction established the functional integrity of the transplanted mitochondria. The MST revealed that Ast-EV-mito transplantation significantly increased basal respiration and ATP production in NMDA-treated primary neurons, but not in the untreated group (Fig. 6H). Collectively, these data indicate that healthy neurons can take up exogenous EV-mito and may not necessarily utilize them for respiration. Alternatively, dysfunctional neurons may use the transplanted exogenous mitochondria for their aerobic respiration and survival, consistent with previous studies in macrophages^65^.

## Discussion

Our study identifies astrocytic mitochondrial transfer as a fundamental mechanism of neuronal protection after traumatic brain injury (TBI), with implications that extend beyond injury to broader principles of glial–neuronal metabolic coupling. Although TBI profoundly alters mitochondrial dynamics in both neurons and astrocytes, the most profound functional deficits occur in neuronal mitochondria, particularly within synaptic compartments that are highly energy-dependent. Astrocytes, in contrast, preserve mitochondrial respiratory capacity and activate transcriptional programs related to organelle trafficking, extracellular vesicle biogenesis, and mitochondrial translation, thereby positioning themselves as effective donors of mitochondria. Consistent with these transcriptional changes, we observed enhanced astrocytic mitochondrial transfer to neurons both *in vivo* following TBI and *in vitro* under excitotoxic conditions. Mechanistically, we further demonstrate that Ast-EV-mito were sufficient to restore mitochondrial function in primary cortical neurons exposed to NMDA, directly linking mitochondrial transfer to neuronal resilience. Together, these findings highlight a previously underappreciated role of astrocytes in dynamically supporting neuronal survival through organelle donation, suggesting that harnessing mitochondrial transfer may represent a therapeutic strategy for neurodegenerative and neurovascular disorders characterized by mitochondrial dysfunction.

### Contrasting neuron and astrocytic mitochondrial bioenergetics following TBI

Currently, brain mitochondrial isolation from our lab and others follows either ficoll/Percoll-based isolation^9,66,67^ or TOM22 antibody-based Fractionated mitochondrial magnetic separation (FMMS)^31^. Both of these methods give rise to two distinct populations of mitochondria, namely, synaptic and non-synaptic. The non-synaptic fraction of mitochondria contains neuronal soma and other glial cell mitochondria, while synaptic mitochondria are isolated from synaptosomes using high-pressure (∼1200 psi) rupture, followed by ficoll/percoll purification. It was impossible to isolate cell-type-specific mitochondria until Dr. Misgeld generated Mito Tag mice, which express a Cre recombinase-dependent GFP protein targeted to the outer mitochondrial membrane (OMM)^68^. Using a GFP antibody, he isolated cell-specific mitochondria from cerebellar Purkinje cells, granule cells, and astrocytes, and determined the mitochondrial proteomic diversity. However, to date, no one has studied cell-type-specific mitochondrial bioenergetics in the brain. In this study, we developed four different mouse models to investigate mitochondrial bioenergetics and dynamics in neurons and astrocytes from sham and TBI mouse models.

Mitochondrial dynamics are strongly associated with mitochondrial bioenergetics ^69–73^. Increased mitochondrial fission is associated with extended physiological demand^74^. However, chronic mitochondrial fission drives pathology and mitochondrial dysfunction^33,34,75,76^. TBI is associated with acute mitochondrial fission^30^. However, cell specificity and associated bioenergetics are not well understood. In our study using NmtD2 and AmtD2 mouse models (expressing Dendra2 from the inner mitochondrial membrane), we confirmed that both neurons and astrocytes respond to injury, exhibiting an increased mitochondrial fission-like phenotype associated with a change in morphology towards a spherical rather than slender shape. To study cell-specific mitochondrial bioenergetics, we used NmtGFP and AmtGFP mouse models (which express GFP from the outer mitochondrial membrane).

From NmtGFP tissues, we isolated four different fractions of mitochondria, namely total, soma, synaptic, and NN , using an anti-GFP antibody. Bioenergetics measurements reveal that out of these four fractions of mitochondria, only synaptic mitochondria were most vulnerable to TBI. This result is consistent with previous studies that used the Ficoll/Percoll method to isolate the synaptic fraction of mitochondria^7,67,77,78^. In our method, we further separated the conventional non-synaptic (neuronal soma+glial) fraction into two fractions, namely, the soma and the NN fraction. Interestingly, we found the least mitochondrial dysfunction from the soma mitochondrial fraction after injury compared to other fractions. Still, the NN fraction (glial-enriched) cells showed significant mitochondrial dysfunction. To study further, we used AmtGFP mice to isolate only astrocyte-specific mitochondria from the glial cell population. Bioenergetics data indicated that astrocytic mitochondria had no injury effect in either the cortex or hippocampus. In contrast, the same samples exhibited significant mitochondrial dysfunction from the total mitochondrial fraction, which includes astrocytic mitochondria. Taken together, though mitochondrial dynamics changed in both neurons and astrocytes, bioenergetics does not seem to have a direct proportional effect at least 24 hrs post-injury. From the conventional non-synaptic portion, we separated astrocytic mitochondrial fractions and found no significant injury effect. Further experiments are in progress with the generation of microglia- and oligodendrocyte-specific mitoGFP mice models to dissect how these cell-type-specific mitochondria are affected by TBI.

### Excitotoxicity and mitochondrial transfer from astrocytes to neurons, both in vivo and in vitro

There is limited *in vivo* evidence supporting the transfer of mitochondria from astrocytes to neurons, with most studies conducted *in vitro*. Mitochondrial transfer between astrocytes and neurons has been shown to occur via CD38/cyclic ADP-ribose (cADPR) signaling and is regulated by mitochondrial Rho GTPases (MIRO1 and MIRO2). This process is disrupted by mutations in GFAP, as seen in Alexander disease, which impairs astrocyte-to-neuron mitochondrial transfer^79^. Activation of astrocytes with cADPR increases the release of extracellular mitochondrial particles into the media, and the addition of astrocyte-conditioned media to neurons subjected to oxygen-glucose deprivation (OGD) improves ATP levels and neuronal viability, suggesting a neuroprotective role for astrocyte-derived mitochondria^20^. In our lab, we were the first to demonstrate intercellular mitochondrial transfer from astrocytes to brain capillaries under physiological conditions using a transgenic mouse model expressing a mitochondria-targeted fluorescent reporter. This transfer was observed in naïve mice that was further enhanced with aging^22^. In a related study, a similar approach was employed using an adeno-associated virus (AAV) system to target astrocytes, showing that low-density lipoprotein receptor-related protein 1 (LRP1) facilitates astrocyte-to-neuron mitochondrial transfer. This occurs by downregulating glucose uptake, glycolysis, and lactate production, thereby reducing ADP-ribosylation factor 1 (ARF1) lactylation and promoting mitochondrial transfer^24^. Collectively, these findings support the existence of astrocyte-to-neuron mitochondrial transfer as a physiologically relevant and regulated process. To date, no studies have directly examined how astrocyte-to-neuron mitochondrial transfer contributes to recovery in overt injury models such as TBI. Consistent with previous findings, we observed that mitochondrial transfer from astrocytes to neurons occurs under physiological conditions, without external stimulation, both in vitro and in vivo (Figures 3 and 6). Notably, this transfer was significantly enhanced following TBI, coinciding with astrocyte activation. It was also increased *in vitro* when neurons were exposed to NMDA, mimicking excitotoxic stress in a neuron–astrocyte co-culture model. Interestingly, our results contrast with a previous study in a mouse stroke model, where inhibition of astrocyte activation using Ginsenoside Rb1 enhanced mitochondrial transfer in vitro. This effect was attributed to reduced mitochondrial complex I activity in astrocytes under conditions of the oxygen–glucose deprivation and reperfusion (OGD/R) model *in vitro*^80^. The discrepancy between studies may stem from differences in experimental design. Specifically, the prior study quantified mitochondrial content indirectly by measuring ATP levels in astrocyte-conditioned media, rather than assessing direct intercellular transfer using co-culture or in vivo models. To our knowledge, this is the first study to demonstrate that TBI enhances mitochondrial transfer from activated astrocytes to neurons *in vivo*. Taken together, this study indicates that brain injury enhances astrocyte-to-neuron mitochondrial transfer, likely mediated by glutamate-induced calcium signaling in astrocytes.

### Activated astrocytes are reprogrammed to generate more TNTs, EVs, and mitochondria

Under physiological conditions, white matter astrocytes and astrocyte end feet surrounding large penetrating vessels exhibit prominent GFAP expression^81^. The functional significance of GFAP is further supported by evidence showing that GFAP deletion impairs reactive astrogliosis^21^ increases mortality following TBI ^82^, and mutations in GFAP associated with Alexander disease impair astrocyte-to-neuron mitochondrial transfer^79^. Both GFAP and vimentin are essential to initiate reactive astrogliosis after TBI to clear debris. However, the precise contribution of GFAP upregulation in activated astrocytes during TBI remains poorly understood. Our results demonstrate that activated astrocytes undergo reprogramming, characterized by altered gene expression associated with mitochondrial homeostasis, mitophagy, TNT formation, EV biogenesis, cargo loading, and secretion (Figure 5). Astrocytes are known to transfer cellular contents to neurons *in vivo* through TNTs or EVs^83–85^. In humans, astrocyte-derived EVs (ADEVs) are increased in plasma one month after stroke ^86^, and also ADEVs isolated from human activated astrocytes have been shown to enhance neuronal uptake, modulate differentiation, and influence neuronal firing^85^. Collectively, these findings suggest that in the acute phase of TBI, astrocytes undergo transcriptional reprogramming to facilitate enhanced intercellular communication and mitochondrial quality control, thereby equipping them to generate, transfer, and eliminate mitochondria through both EV- and TNT-mediated mechanisms.

### Therapeutic feasibility of Ast-EV-mito transfer/transplantation for excitotoxicity

Recently, EMT has emerged as a promising therapeutic strategy for diseases characterized by mitochondrial dysfunction, including cardiovascular, respiratory, and neurodegenerative disorders^87–90^. Transfer of healthy mitochondria appears to be an effective rejuvenation process in damaged cells ^90,91^. Previous studies have shown that EMT using EV-mito improves neuronal ATP levels and viability *in vitro*^20^. However, it remains unknown whether EV-mito can mitigate excitotoxicity-induced neuronal dysfunction, a common pathological mechanism underlying several neurodegenerative diseases, including TBI.

In this study, we established that NMDA-mediated excitotoxicity significantly impairs mitochondrial function and alters mitochondrial dynamics in neurons (Figure 7). Remarkably, we were able to rescue this mitochondrial dysfunction by transplanting healthy mitochondria isolated from astrocyte-conditioned media. These mitochondria, Ast-EV-mito, were endogenously labeled with a mitochondrial-targeted fluorescent reporter (mtD2), which eliminates concerns associated with nonspecific signals due to dye leakage, a common limitation in mitochondrial transplantation studies using exogenous dyes, such as MitoTracker. Furthermore, using high-resolution confocal microscopy coupled with Imaris 3D reconstruction, we demonstrated that transplanted mitochondria not only internalize into recipient neurons but also colocalize with the endogenous mitochondrial marker TOM20, confirming successful integration rather than surface adherence. To directly assess the functional impact of mitochondrial transplantation, we measured neuronal bioenergetics using the Seahorse XF flux analyzer, thereby avoiding the limitations of indirect colorimetric ATP and cell viability assays. Taken together, this study is the first to demonstrate the therapeutic potential of Ast-EV-mito in restoring neuronal mitochondrial function under excitotoxic conditions, a pathophysiological mechanism common to neurodegenerative diseases, including TBI.

### Limitations of the study

While this study provides novel insights into astrocyte-mediated mitochondrial transfer and its potential therapeutic implications following TBI, several limitations should be acknowledged. First, the analysis was conducted at a single post-injury time point, which may not capture the full temporal dynamics of mitochondrial transfer and recovery processes. Second, although the study focused on astrocyte-derived mitochondria, contributions from other glial or neuronal sources cannot be completely excluded. Similarly, the mitochondrial transfer to cell types other than neurons. Third, while *in vitro* and *in vivo* data support the functional integration of transferred mitochondria, additional studies using high-resolution live imaging and genetic labeling are needed to confirm long-term mitochondrial incorporation and bioenergetic impact in recipient neurons. Furthermore, the experimental model may not fully replicate the complexity of human TBI pathology, and extrapolation to clinical settings should be approached with caution. Finally, while mitochondrial transplantation shows therapeutic promise, the precise mechanisms regulating mitochondrial uptake, trafficking, and sustained function in injured neurons remain to be fully elucidated.

## Conclusion

Our findings demonstrate that TBI induces a shift in astrocyte phenotype that promotes mitochondrial packaging and transfer to other cells. Astrocytic mitochondrial transfer increased mitochondrial transfer to the neuronal soma but not to synaptic compartments after TBI. This compartment-specific pattern of transfer aligns with our functional data: neuronal soma receiving astrocytic mitochondria maintained mitochondrial bioenergetics and preserved respiratory complex protein levels, whereas synapses, where no increase in astrocytic mitochondrial delivery was detected, exhibited pronounced mitochondrial dysfunction after TBI. These results suggest that insufficient mitochondrial support at synapses contributes to excitotoxic injury and impaired synaptic metabolism. Whether intrinsic proteomic changes within synaptic or somatic mitochondrial populations underlie this differential vulnerability remains an important question for future investigation. Importantly, we observed that astrocytes can release EV-mito, and our proof-of-concept *in vitro* experiments demonstrate that EV-mito supplementation enhances mitochondrial transfer and improves neuronal bioenergetics in an NMDA-induced excitotoxicity model. Together, these findings identify EV-mito as a promising therapeutic strategy to specifically target synaptic mitochondrial deficits that arise following TBI and excitotoxic signaling.

### Author Contributions

G.V.V., H.J.V., and P.G.S. developed the concept and designed the research. G.V.V. and H.J.V. performed the experiments. G.V.V. analyzed the data, interpreted the experimental results, and prepared the figures. J.M.M. and K.S. did the scRNAseq and analysis. G.V.V., J.M.M., and W.B.H. drafted the manuscript. P.G.S. supervised all the experiments. G.V.V., H.J.V., A.G.R., J.M.M., S.P., W.B.H., and P.G.S. contributed to the editing and critical revision of the manuscript. P.G.S. approved the final version of the manuscript. All authors have read and agreed to the published version of the manuscript.

### Funding

Funding was provided through the VA Merit Award 2I01BX003405 (PGS); National Institutes of Health National Institute of Neurological Disorders and Stroke 1R01NS112693-01A1 (PGS); Kentucky Spinal Cord and Head Injury Research Trust (KSCHIRT), grant 20-7A (PGS); grant 24-14 (PGS and GVV); grant 24-8 (WBH and GVV); NIH (P20 GM148326) (PGS); RF1NS118558 and R01AG070830 (JMM). The contents do not represent the views of the U.S. Department of Veterans Affairs or the United States Government. The content is solely the responsibility of the authors and does not necessarily represent the official views of the NIH.

### Data Availability Statement

The original contributions presented in the study are included in the article/supplementary material. Further inquiries can be directed to the corresponding author.

## Acknowledgments

We would like to thank Frances Meredith and Erin Sullivan (University of Kentucky) for their invaluable assistance with surgical procedures, animal breeding, and colony maintenance. We are grateful to the Spinal Cord and Brain Injury Research Center and the Light Microscopy Core at the University of Kentucky for providing access to confocal microscopy and Imaris software for imaging and data processing. We also thank the Mitochondrial Bioenergetics Core for access to Seahorse XF analysis.

## Conflicts of Interest

The authors declare no conflict of interest.

## Institutional Review Board Statement

All the studies performed were approved by the University of Kentucky IACUC, which the Association accredits for the Assessment and Accreditation for Laboratory Animal Care, International (AAALAC, International), and all experiments were performed in accordance with its guidelines. PHS Assurance #D16-00217 (A3336-01); Protocol #2021-3849.

## Supplementary information

**Figure S1:**
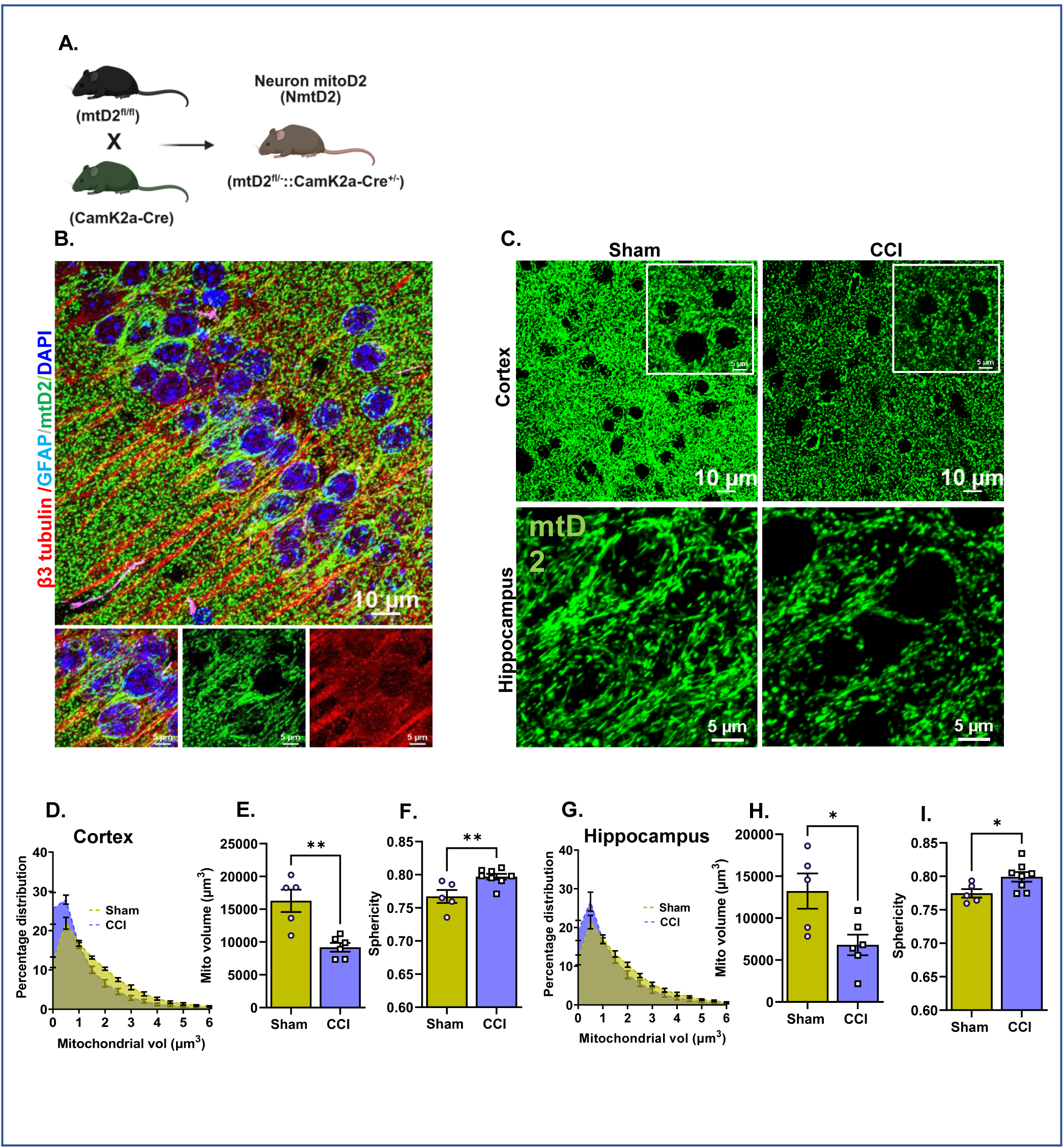
TBI alters neuronal mitochondrial dynamics. **(A)** Schematic illustrating the generation of neuron-specific mitochondrial reporter mice (mtD2^f/f^; CamK2a-Cre^+/−^) expressing the green fluorescent protein Dendra2 (NmtD2) targeted to the inner mitochondrial membrane. (B) Representative confocal micrograph of a hippocampal brain section from NmtD2 mice, stained for β3-tubulin (red), glial fibrillary acidic protein (GFAP-pink), and DAPI (blue). (C) Representative micrographs of brain sections from NmtD2 mice 24 hrs post-CCI or sham surgery (D, E) Quantification of mitochondrial morphology (green) using Imaris software from the ipsilateral penumbral cortex and hippocampus. The histograms display the percentage distribution of mitochondrial volume while the bar graphs illustrate the total mitochondrial volume and sphericity. Each circle/square represents one animal. Data represented as Mean ± SEM. P ≤ 0.05 *; P ≤ 0.01** by unpaired t-test.

**Figure S2:**
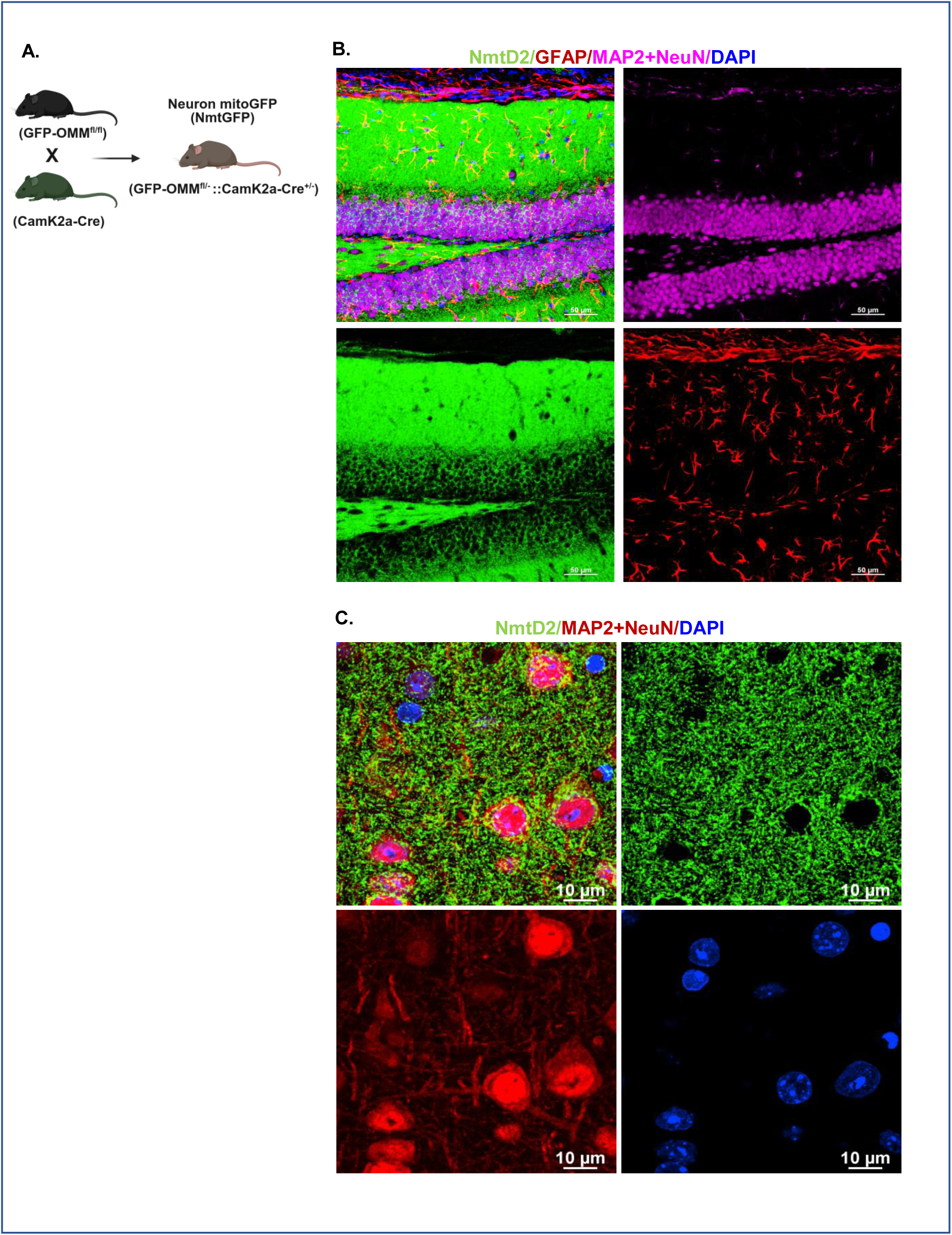
Generation of neuron-specific mitochondrial GFP mice (NmtGFP): (A) Breeding scheme of neuron-specific mitochondrial reporter mice (NmtGFP). (B, C). Representative confocal micrograph of NmtGFP mice brain section, immunostained with GFAP (pink) and MAP2+NeuN (pink), nucleus (blue), and transgene mtGFP (green).

**Figure S3:**
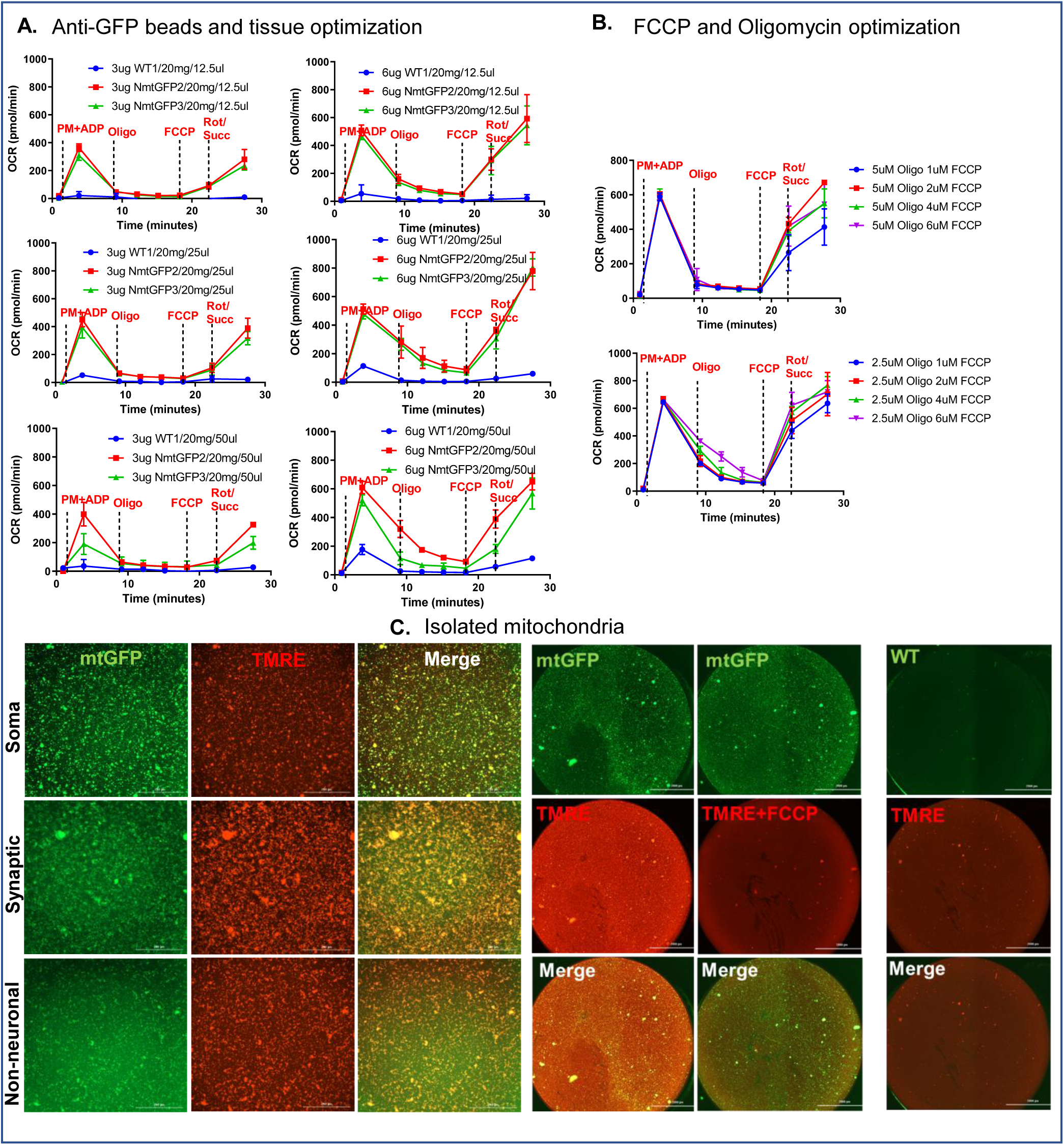
Optimization of cell-specific mitochondrial isolation Bioenergetics study: **(A)** OCR traces from the neuronal fraction of mitochondria. The tissue amount (grams) was narrowed down based on our published work and pilot experiments. Approximately 1 µl of anti-GFP antibody for 1mg or 0.5mg of tissue was used in both WT and NmtGFP brain homogenates to verify the specificity of binding. Two different concentrations of mitochondria (3 µg and 6 µg) were loaded into each SeahorseXFe96 well plate to check the saturation point. (B) OCR traces from neuronal fraction of mitochondria optimized for FCCP and Oligomycin concentration. 2.5 and 5 µm Oligomycin tested with 1, 2, 4, and 6 µM of FCCP concentration. (C) Representative images of anti-GFP bead isolated mitochondria (green) from NmtGFP and WT mice (no mitoGFP tag). Isolated mitochondria were stained with TMRE (red) to test whether isolated mitochondria maintain intact membrane potential. Some wells were pretreated with FCCP to disrupt membrane potential before staining with TMRE (No TMRE (red) stain).

**Figure S4:**
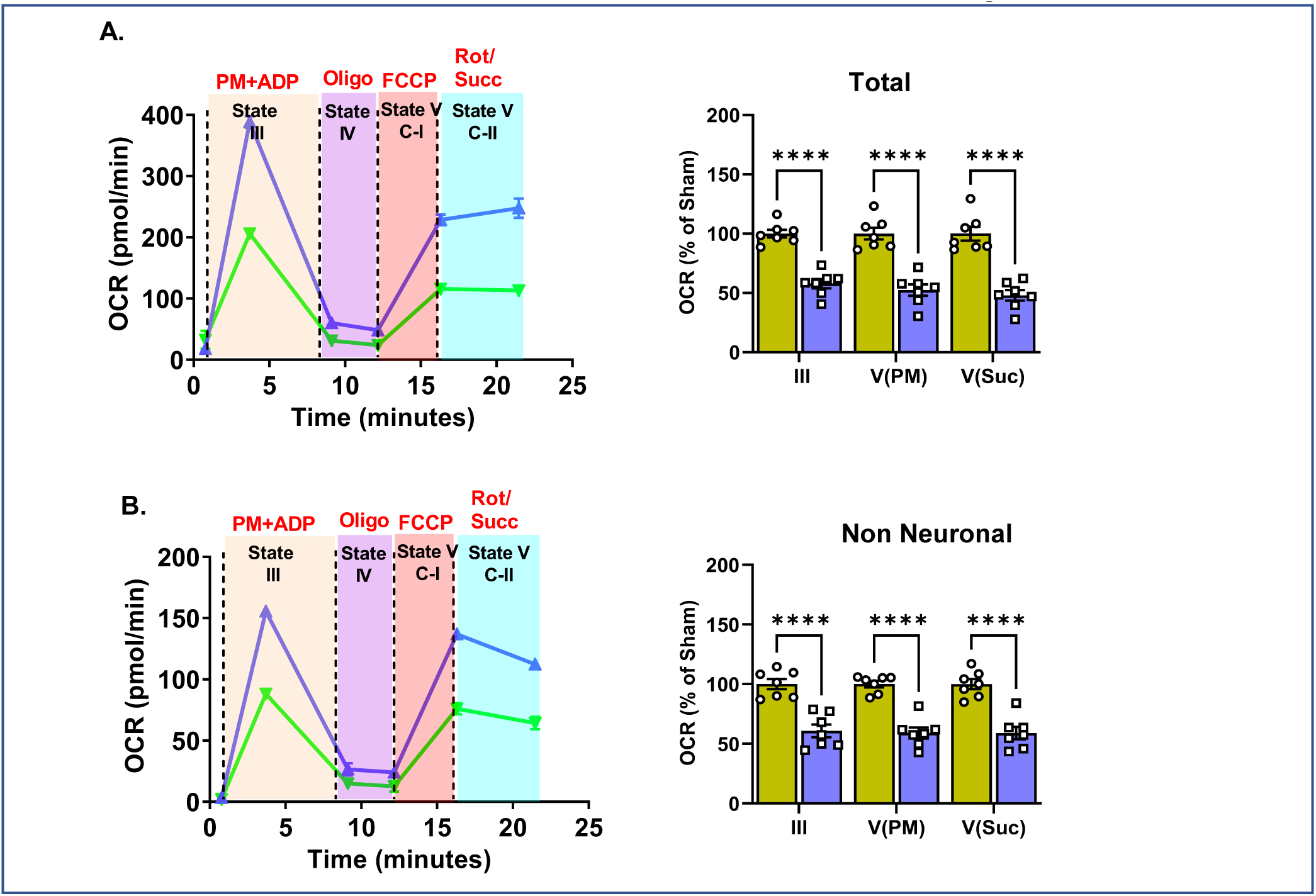
Total and NN fraction of mitochondrial bioenergetics. (A, B, C, D) Representative traces and quantification of OCR from total and non-neuronal mitochondria isolated from sham and CCI (ipsilateral punch). Each circle/square represents one animal (n = 7 mice/group).

**Figure S5:**
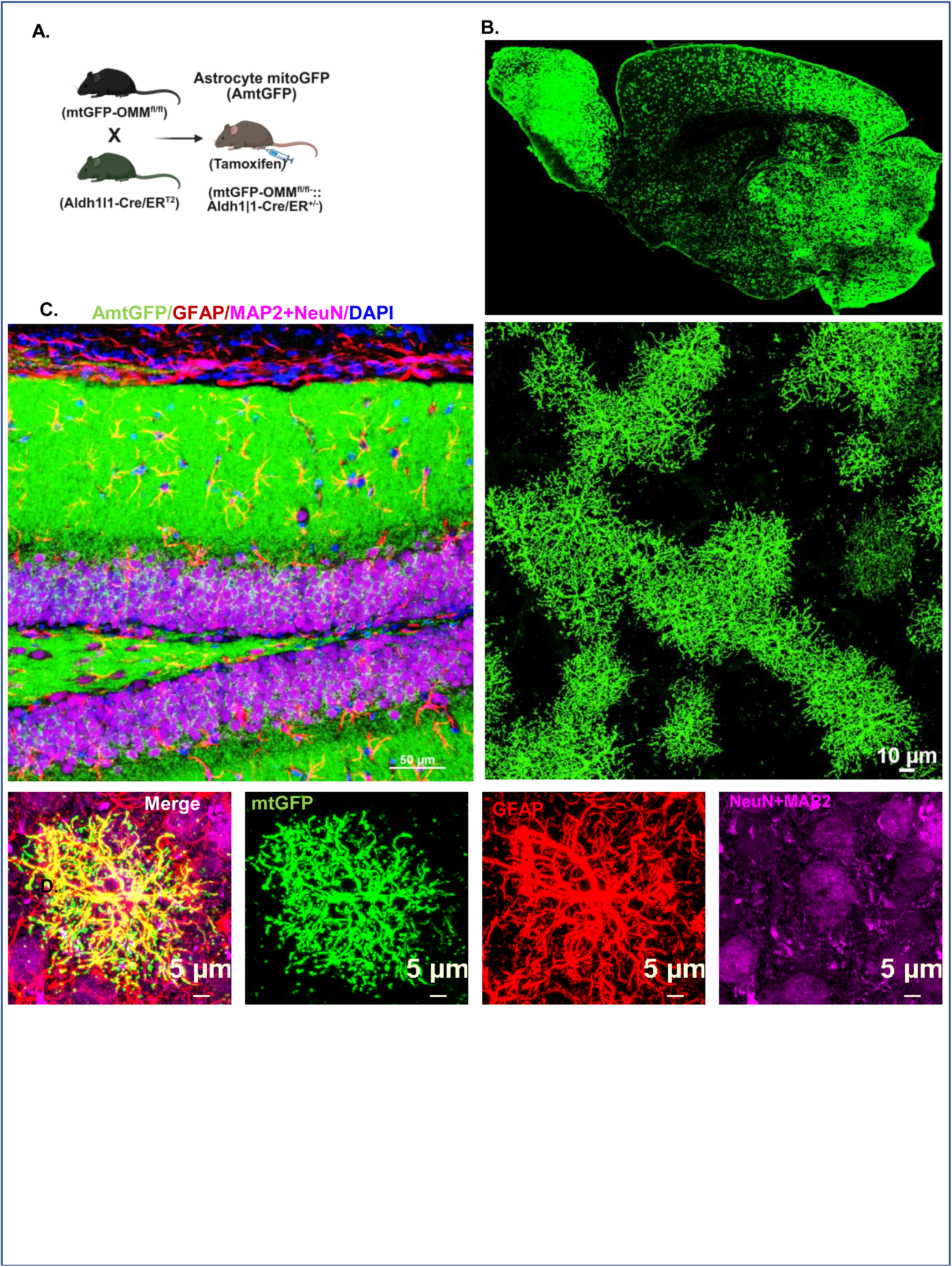
Generation of Astrocyte-specific mitochondrial GFP mice (AmtGFP): A) Breeding scheme of Astrocyte-specific mitochondrial reporter mice (AmtGFP). (B, C). Representative confocal micrograph of NmtGFP mice brain section. (D). Representative confocal micrograph; GFAP (red) and MAP2+NeuN (pink), nucleus (blue), and transgene mtGFP (green).

**Figure S6:**
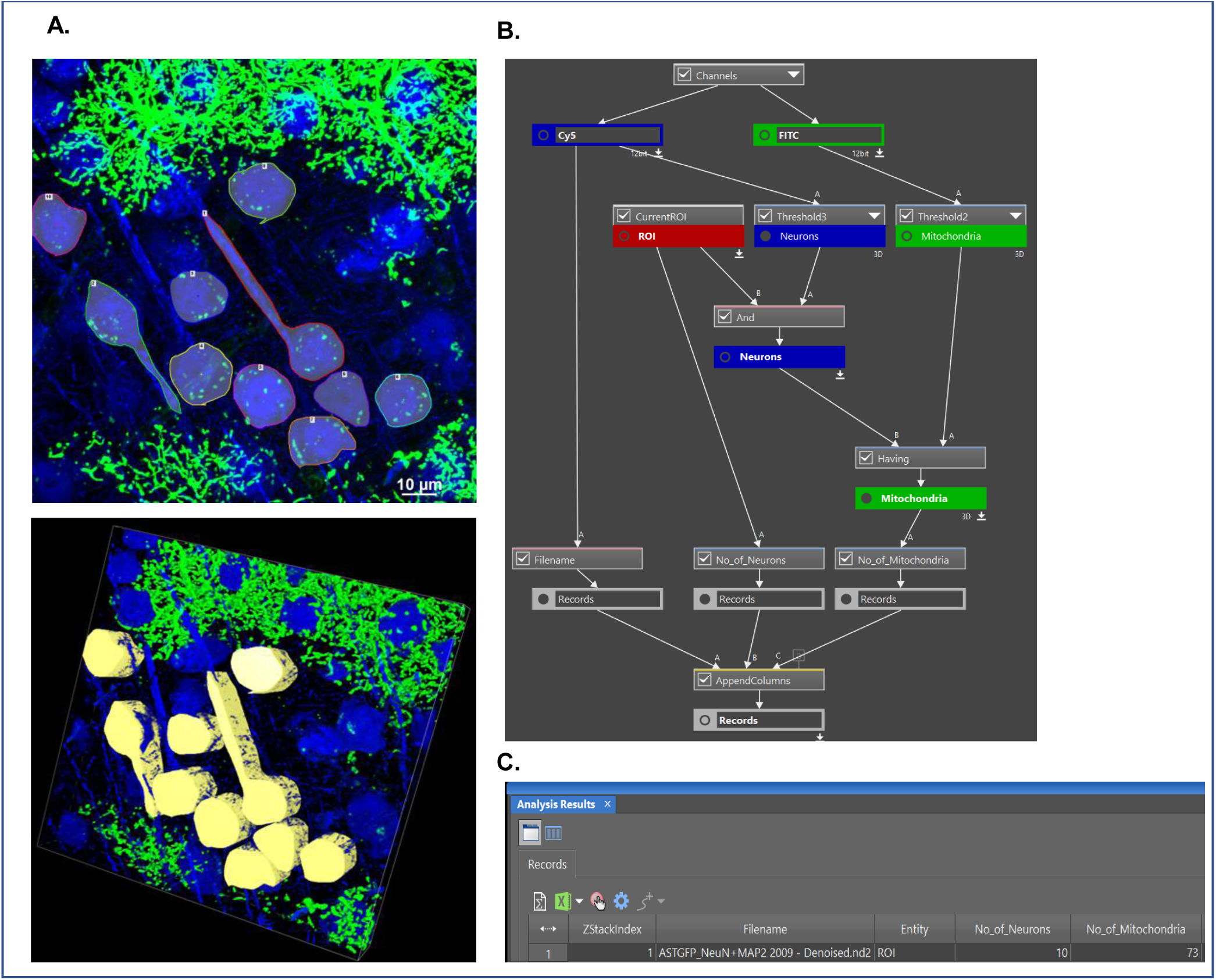
Unbiased astrocyte derived mitochondrial count in neuron using Nikon NIS-elements general analysis software. (A) Representative confocal z-stack and 3D images with selected region of interest (ROI) traces for the quantification of astrocytic mitochondrial transfer to neurons. (B, C) The workflow in NIS-Elements general analysis quantifies the number of neurons and mitochondria within the ROI in a 3D image.

## Notes

### Competing Interest Statement

The authors have declared no competing interest.

